# Overexpression of Eaf1, a subunit of the NuA4 lysine acetyltransferase complex, rescues growth defects in the *Saccharomyces cerevisiae* H3K36M oncohistone model via histone H4 tail acetylation

**DOI:** 10.64898/2026.05.30.728992

**Authors:** Celina Y. Jones, Yu-Hsiang Laurence Chen, Milo B. Fasken, Anita H. Corbett, Jennifer M. Spangle

## Abstract

Histone proteins are critical for regulating nucleic acid transactions, including gene expression. Specific missense mutations in histone genes have been linked to oncogenesis, creating oncohistones. One of the first identified oncohistones converts K36 in histone H3 to M (H3K36M). While humans have 15 H3 genes, the *Saccharomyces cerevisiae* genome contains only two H3 genes. As H3 shares 90% sequence identity between budding yeast and humans, the budding yeast system provides an ideal model for study of H3K36M. We previously identified the catalytic subunit of the NuA4 lysine acetyltransferase complex as a high copy suppressor of growth defects in H3K36 mutant yeast cells. Here, we demonstrate that the NuA4 scaffold Eaf1 suppresses mutant H3K36 growth defects. We show that this suppression by Eaf1 is dependent on the Eaf1 HSA domain. To examine whether the role of NuA4 in acetylating histone H4 is required for suppression of H3K36 growth defects, we altered the four NuA4 target lysines within H4 (K5, 8, 12, and 16) to create quadruple histone mutants with all four of these lysines converted to alanine (H4-4K→A) or arginine (H4-4K→R). These H4 mutants have reduced viability and disrupted global H4 acetylation. Eaf1 cannot suppress growth defects in these H4 acetylation mutants, suggesting Eaf1-mediated suppression is dependent on acetylation of H4 N-terminal tails. We further show that inhibiting NuA4 activity in H3K36M human cells limits their oncogenic potential. Together, this work defines a mechanistic connection between H3K36 and H4 N-terminal tail acetylation in the context of a cancer-causing mutation.

**Article Summary:** Specific mutations in histone genes have been found to cause various cancers, creating oncohistones, and the exact mechanisms by which they drive oncogenesis remain to be characterized. Budding yeast is a simple yet powerful model for studying oncohistones such as H3K36M. Subunits of the NuA4 lysine acetyltransferase complex were found to suppress growth defects in budding yeast expressing H3K36M. This suppression was dependent on the presence of the NuA4 target lysines in histone H4. Additionally, inhibiting NuA4 activity in human H3K36M cells limited their oncogenic potential. This work characterizes the mechanistic relationship between H3K36M and the NuA4 complex.

## Introduction

Histone proteins are crucial for compacting DNA and regulating gene expression. Genomic DNA is packaged by wrapping around histone heterooctamers, creating the basic repeating nucleosome structure (Luger et al. 1997). The accessibility of other proteins that act upon DNA is controlled by local DNA coiling and compaction of nucleosomes. DNA accessibility impacts various key genomic functions, such as DNA repair and gene expression (Mitchener and Muir 2022). Chromatin accessibility and recruitment of chromatin modifying enzymes is regulated by post translational modifications (PTMs) on histones. There are many genes encoding histone proteins in humans, from 12-15 genes per histone H2A, H2B, H3, and H4 present (Zhang et al. 2023). Therefore, the discovery that missense mutations in a single allele of an individual histone gene could cause cancers despite this genetic redundancy was surprising (Mitchener and Muir 2022). One of the first histone mutants identified that drives oncogenesis is histone H3 with lysine converted to methionine at residue 36 (H3K36M) (Behjati et al. 2013; Papillon-Cavanagh et al. 2017). Such histone proteins that confer oncogenic driver activity are termed oncohistones. H3K36M drives head and neck squamous cell carcinomas and chondroblastomas and has also been implicated in some other cancers (Behjati et al. 2013; Papillon-Cavanagh et al. 2017).

The primary mechanism by which characterized oncohistones, including H3K36M, drive oncogenesis is due to the disruption of crucial histone PTMs that are associated with transcriptional regulation (Lewis et al. 2013; Lu et al. 2016). Methylation at H3K36 is tightly controlled due to a key role in regulating gene expression (Wagner and Carpenter 2012). Critically, cells expressing the oncohistone H3K36M show reduced global methylation at H3K36me2/3 (Lu et al. 2016; Zhang et al. 2017). The H3K36me3 methyltransferase SETD2 binds H3K36M with higher affinity than native H3K36, trapping SETD2 and inhibiting deposition of methyl groups on H3 proteins both in *cis* and in *trans* (Zhang et al. 2017; Lu et al. 2016). This *cis* and *trans* regulation renders a single oncohistone mutant protein capable of dysregulating nucleosome-associated wild type (WT) histone H3 proteins in a dominant negative manner. A similar mechanism has been reported for cancers characterized by other oncohistone mutations, including gliomas caused by H3K27M mutations (Justin et al. 2016).

Studying the oncohistone H3K36M in the context of WT histone H3 expression is important to define mechanisms underlying oncogenesis. However, examining the biological consequences of the H3K36M amino acid change is challenging in a system with so many copies of histone genes. Such mechanistic studies can be streamlined and simplified by exploiting genetic models. The high number of histone genes in mammalian models including human cells presents a technical challenge for engineering cell lines that express only the oncohistone protein, but the genome of the model system *Saccharomyces cerevisiae* encodes only two histone H3 genes, *HHT1* and *HHT2*, compared to the human genome which encodes 15 histone H3 genes. Thus, the oncogenic mutation under study can be readily engineered at one H3 gene in budding yeast and the other H3 gene deleted so that only the oncohistone protein is expressed. Furthermore, human and budding yeast histone H3 proteins share 90% sequence identity (Zhang et al. 2023), making *S. cerevisiae* a valuable model to characterize the functional biology of oncohistone mutants.

Our group previously used a budding yeast H3K36-mutant model in a high copy suppressor screen to rapidly identify novel suppressors of oncohistone-driven growth defects (Lemon et al. 2022). One of the suppressors that we identified was Esa1, the catalytic component of the NuA4 lysine acetyltransferase complex, which is conserved in mammals as the TIP60 complex (Lemon et al. 2022). While numerous NuA4/TIP60 histone and non-histone substrates have been defined, a predominant role of NuaA4 is the acetylation of H4 and H2A tails (Downey et al. 2015; Lin et al. 2009; Yi et al. 2012; Allard et al. 1999). In the initial study, we demonstrated that Esa1-mediated suppression was dependent on the catalytic acetyltransferase function (Lemon et al. 2022), but the specific mechanism was not further defined (Downey et al. 2015; Lin et al. 2009; Yi et al. 2012; Allard et al. 1999).

NuA4 complex-driven acetylation is crucial for regulating processes such as transcription and DNA repair (Rossetto et al. 2014; Jacquet et al. 2016; Squatrito et al. 2006). There are 13 subunits in the NuA4 complex, and 12 of these subunits are conserved between budding yeast and humans (Doyon et al. 2004; Qu et al. 2022). An independent subcomplex of NuA4, called piccolo NuA4 (picNuA4), is comprised of the subunits Esa1, Eaf6, Epl1, and Yng2 (Boudreault et al. 2003; Friis et al. 2009). The full NuA4 complex is built around the scaffolding subunit Eaf1 (Wang et al. 2018). Both the complete NuA4 and picNuA4 complexes are depicted in Fig. 1A with subunits relevant to this study labeled. PicNuA4 and the full NuA4 complex have different preferential binding along gene bodies, where NuA4 binds more tightly at promoters, implicating the full complex in transcription initiation, and picNuA4 binds equally well along the coding region, implicating the subcomplex in transcription elongation (Ginsburg et al. 2009). In both budding yeast and humans, methylation at histone H3 helps recruit NuA4 to nucleosomes. Specifically, H3K4me1/2/3 and H3K36me2/3 direct the NuA4 complex to nucleosomes via the Esa1 and Eaf3 chromodomains and enhances its lysine acetyltransferase activity (Ginsburg et al. 2014; Su et al. 2016; Li and Wang 2017; Steunou et al. 2016). This connection between H3 methylation and H4 acetylation suggests an explanation for why Esa1 was identified as a suppressor of growth defects in our prior H3K36-mutant screen. Reduced H3 methylation in budding yeast exclusively expressing the H3K36 mutant may decrease recruitment of NuA4 to nucleosomes. Thus, the overexpression of the NuA4 Esa1 subunit may overcome this decrease in recruitment.

**Figure 1.**
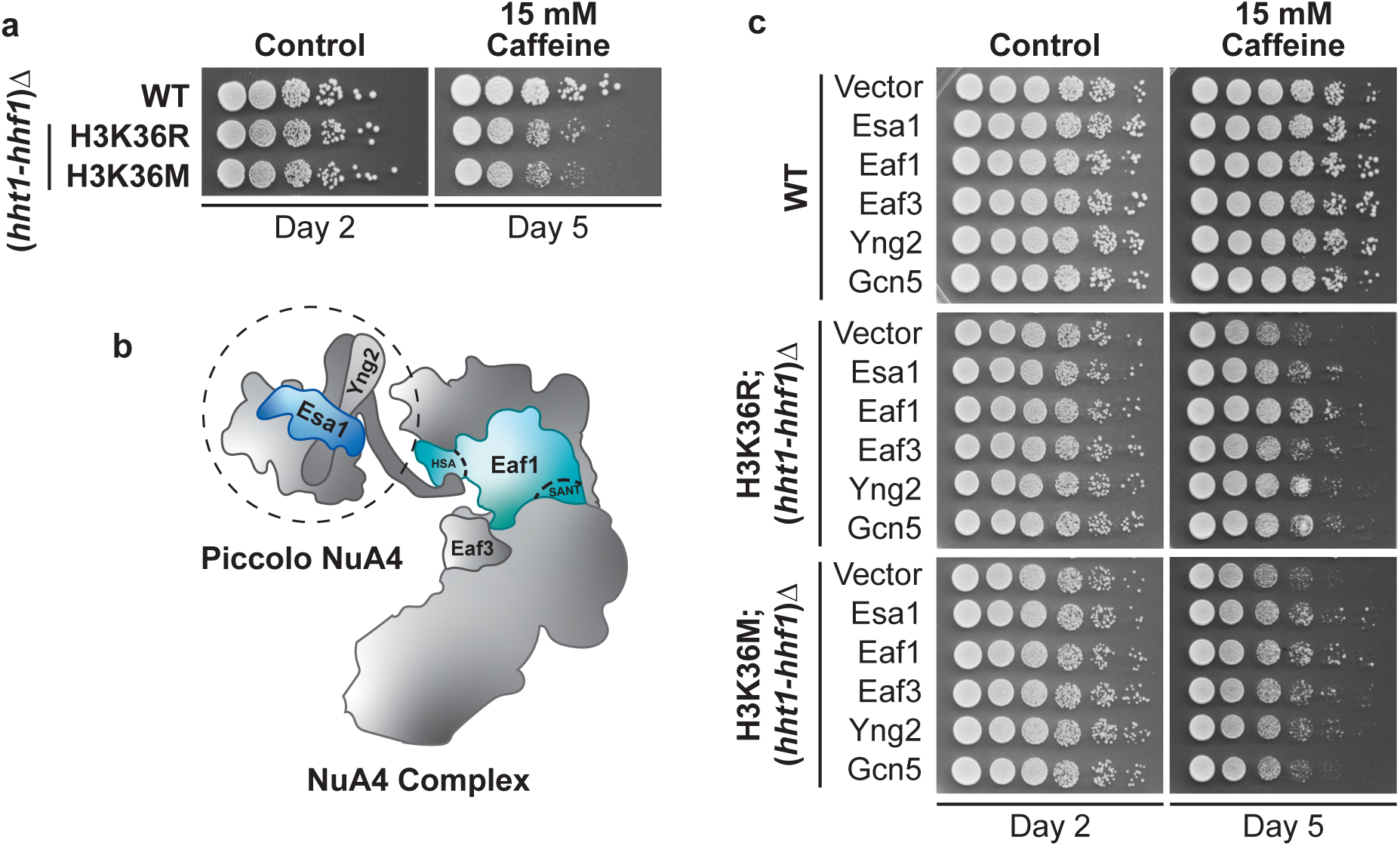
Esa1 and Eaf1 suppress growth defects in H3K36 mutant cells. a) A diagram of the NuA4 complex, with Esa1 in blue and Eaf1 (and its domains HSA and SANT) in teal. Yng2 and Eaf3 are also labeled in gray. The other nine subunits not investigated in this study are unlabeled. The piccolo NuA4 subcomplex, that is independently catalytically active, is marked by a dotted circle. b) Cells expressing H3K36R or M as the sole copy of histone H3 display growth defects when grown on plates containing Caffeine in a serial dilution growth assay but not on Control plates. c) The ability of a subset of NuA4 components to rescue growth defects in H3K36R/M cells when overexpressed on high copy vectors was assessed. Growth was assessed on both Control plates and plates containing Caffeine. Both Vector alone and Gcn5, the catalytic component of the SAGA complex, were included as controls. Alt text: a) Yeast growth assays comparing the growth of H3K36R and H3K36M cells on caffeine to that of WT. b) A diagram of the NuA4 complex, with the relevant subunits labeled. c) Yeast growth assays displaying the ability of various NuA4 subunits to suppress growth defects in WT, H3K36R, and H3K36M cells.

Here, we sought to characterize the mechanistic basis of NuA4-mediated suppression of growth defects in H3K36M budding yeast cells. We identified the NuA4 subunit Eaf1, the complex scaffold, as a stronger suppressor of growth defects than Esa1 and show that suppression by Eaf1 is dependent on the presence of the Eaf1 HSA protein domain, which is necessary to maintain complex integrity. We focused on NuA4 target substrates on histone H4 N-terminal tails, creating mutants that eliminate NuA4-mediated H4 acetylation. These H4 mutants have reduced viability and disrupted H4 acetylation. Importantly, neither Esa1 nor Eaf1 suppresses H3K36 mutant growth defects when H4 acetylation is genetically eliminated. Finally, we translated these findings to a human H3K36M cell line and provide evidence that inhibiting NuA4 function reduces oncogenic potential in these H3K36M oncohistone-expressing cells. This work further defines the functional connection between histones H3K36 and H4 N-terminal tails and serves as a template for rapid characterization of cancer-causing human mutations in a budding yeast system.

## Materials and Methods

### *S. cerevisiae* strains and chemicals

Chemicals used for experiments with *Saccharomyces cerevisiae* were obtained from Sigma-Aldrich (St. Louis, MO), United States Biological (Swampscott, MA), or Fisher Scientific (Pittsburgh, PA) unless otherwise noted. All media were prepared by standard procedures (Adams et al.). All DNA manipulations were performed according to standard procedures (Sambrook et al. 1989). *S. cerevisiae* strains and plasmids used in this study are listed in Table S1. The PCR- and homologous recombination-based system for generating targeted mutations in histone genes in budding yeast cells has been described (Duina and Turkal 2017). Histone H3 mutations at the endogenous *HHT2* gene were generated using the parental *hht2Δ::URA3:TRP1* strain (yAAD165) and the strategy detailed previously (Johnson et al. 2015; Duina and Turkal 2017). The endogenous *HHT1* and *HHF1* genes in these strains were subsequently deleted and replaced via homologous recombination with a *kan*MX marker cassette. Histone H4 mutations at the endogenous *HHF2* gene were generated in these strains utilizing a similar method to that in (Johnson et al. 2015; Duina and Turkal 2017). However, the method differed in that the strains already harbored a deletion of *HHT1-HHF1*. A pRS315 plasmid expressing *HHT2-HHF2* was co-transformed into the cells to enable successful integration of the H4 mutants. This plasmid was retained in the strains for subsequent experiments by maintaining them in dropout media.

### Plasmids

Plasmids expressing *ESA1*, *EAF1*, *EAF3*, *YNG2*, or *GCN5* were cloned into the YEp352 vector (Hill et al. 1986). The Eaf1 domain mutants (eaf1-HSAΔ, eaf1-SANTΔ, eaf1-HSAΔSANTΔ) in a pRS316 vector (Sikorski and Hieter 1989) were generously provided by Michael Kobor’s group (Lu et al. 2022). The WT *HHT2-HHF2* maintenance plasmids were on either pRS315 or pRS316 vectors (Sikorski and Hieter 1989).

### *S. cerevisiae* serial dilution growth assays

To examine the growth of mutant cell lines, yeast were grown overnight at 30°C to saturation in 2 mL YEPD (yeast extract, peptone, dextrose) media. Cells were normalized to OD_600_ = 5, serially diluted in 10-fold dilutions, spotted on control YEPD media plates or YEPD media plates containing 15 mM caffeine and grown at 30°C for 2-5 days. Cells were also grown on Ura- plates containing 2% glucose to maintain plasmid. To test the effect of suppressor plasmids on the growth of mutant H3 and H4 cells in the presence of caffeine, cells were transformed with plasmid as noted in the respective figure and detailed in Table S1. Cells were then grown overnight at 30°C to saturation in 2 mL Ura- or Leu- media containing 2% glucose. Cells were normalized to OD_600_ = 5 and serially diluted as previously described, spotted onto control YEPD media plates or YEPD media plates containing 15 mM caffeine, and grown at 30°C for 2-5 days. Cells were also spotted onto control Ura- or Leu- plates.

### Double histone mutant viability assay

To investigate whether H4 mutants (H4-4K→A or H4-4K→R) alone or combined with H3 mutants (H3K36R or H3K36M) are viable, we assessed the viability of each mutant in the absence of WT histone H3 or H4 expression. Cell lines were established as described above, and each expressed a pRS316 plasmid containing WT *HHT2-HHF2*. Cells were grown overnight at 30°C to saturation in 2 mL Ura- media, and cell concentration was estimated with OD_660_ reads. Cells were then diluted and plated at 200 cells/10-cm dish on Ura- and 5-fluoroorotic acid (5-FOA) plates. Cells grew for 3-7 days, plates were imaged, and colonies were counted. The data were graphed in GraphPad Prism 11 (GraphPad Software, LLC) as a ratio of the number of colonies that grew on 5-FOA plates normalized to the number of colonies on the respective Ura- plates with standard deviation (n=4). Statistical significance was assessed via one-way ANOVA analyses.

### Mammalian cell lines

Human cell lines were generated and maintained as described in (Sad et al. 2025). Briefly, wildtype *D. melanogaster* H3.3 (NCBI NM_001273153), which shares 100% amino acid identity with *H. sapiens H3.3,* was cloned into the pBabePuro IRES GFP plasmid (Addgene #14430). The H3.3K36M mutation was introduced into the pBabePuro dH3.3-IRES GFP using site-directed mutagenesis (Agilent). Plasmids were verified by sequencing, packaged into retroviral particles, and used to generate stable HMEC cell lines. Authenticated HMEC cells were purchased from ATCC, engineered to express dominant negative p53 (Zhao et al. 2005) and used for experiments. HMEC cultures were maintained at 37°C and 5% CO_2_ in DMEM/F12 media supplemented with 0.6% FBS, 0.01 µg/ml EGF, 10 µg/ml insulin, 0.025 µg/ml hydrocortisone, 1 ng/ml cholera toxin, 2.5 µg/ml Amphotericin B, and 1% Penicillin-Streptomycin.

### Yeast and mammalian cell acid extraction and immunoblotting

For yeast experiments, the indicated yeast cells were grown overnight at 30°C to saturation in 5 mL YEPD (yeast extract, peptone, dextrose) or dropout (Ura- or Leu-, when expressing plasmid) media. Cells were diluted in 100 mL medium and grown at 30°C to a final OD_600_ = 1.0. Cells were pelleted by centrifugation at 1,962 x *g* in 50 mL tubes, washed in ddH_2_O, transferred to 2 mL screwcap tubes, washed again in ddH_2_O, and pelleted by centrifugation at 900 x *g*. Pelleted cells were washed in Buffer 1 (1 M sorbitol, 50 mM Tris-HCl, pH 7.5, 5 mM MgCl_2_) and weighed. Samples were separated into two tubes to be <100 mg each. Cells were resuspended in 1 mL Buffer 1, and 5.4 μL 2-mercaptoethanol (14.3 M) was added to each. Samples were incubated on ice for 10 min. Cells were pelleted at 4°C at 900 x *g* and resuspended in 1 mL Buffer 1 plus 600 μg zymolyase 20T, then incubated at 35°C for 20 min with gentle agitation. Then, 0.7 mL Buffer 2 (1 M sorbitol, 50 mM MES, pH 6, 5 mM MgCl_2_) plus protease inhibitors (Thermo Scientific A32955) was added, and cells were pelleted at 1233 x *g* in 4°C. Samples were resuspended in 0.7 mL Buffer 3 (50 mM MES, pH 6, 75 mM KCl, 0.5 mM CaCl_2_, 0.1% NP-40) plus protease inhibitors, incubated on ice for 5 min, and pelleted at 12,175 x *g* at 4°C. Samples were resuspended in 0.7 mL Buffer 4 (10 mM MES pH=6, 430 mM NaCl) plus protease inhibitors and 0.5% IGEPAL CA-630, incubated on ice for 5 min, and pelleted at 14,489 x *g*. Finally, samples were resuspended in 0.7 mL Buffer 4 plus protease inhibitors, incubated on ice for 5 min, and pelleted at 17,005 x *g*. In brief, pellets were acid extracted in 120 μL 0.25 M HCl and spun in a rotor wheel at 4°C for at least 2 hr. Samples were pelleted at 14,489 x *g* in 4°C, and the supernatants were combined with 8 volumes of acetone and incubated overnight at -20°C. Extracts were pelleted at room temperature at 1,962 x *g*, resuspended in acidified acetone (120 mM HCl in acetone), and pelleted 12,300 x *g*. Extracts were washed in acetone and pelleted. Extracted histones were air dried and resuspended in 50 μL ddH_2_O plus 1 μL NaOH. Protein lysate concentration was determined by Pierce BCA Protein Assay Kit (Life Technologies). Protein lysate samples (20-25 µg) in reducing sample buffer (50 mM Tris HCl, pH 6.8; 100 mM DTT; 2% SDS; 0.1% Bromophenol Blue; 10% Glycerol) were resolved on 4–20% Criterion™ TGX Stain-Free™ precast polyacrylamide gels (Bio-Rad). Protein lysate samples were transferred to nitrocellulose membranes (Bio-Rad) in Dunn Carbonate Buffer (10 mM NaHCO_3_, 3mM Na_2_CO_3_, pH 9.9, 20% methanol) at 22V for 90 min at room temperature, and the resulting membranes were blocked in TBST + 5% milk. After primary and secondary antibody incubation, proteins of interest were visualized and quantified using the ChemiDoc MP Imaging System (Bio-Rad).

Histones were acid extracted from human cells as described previously (Spangle et al. 2016; Jones et al. 2022). In brief, cells were lysed on ice in Triton extraction buffer (TEB; PBS, 0.5% Triton X-100) supplemented with protease inhibitors [7.5 μM aprotinin, 0.5 mM leupeptin, 250 μM bestatin, 25 mM AEBSF-HCl]. Cell lysates were centrifuged at 6,500 x g at 4°C and histones were acid extracted from the resulting pellet with 1:1 TEB:0.8 M HCl. Histones were centrifuged at 4°C and the supernatant was precipitated with the addition of an equal volume of 50% TCA, then centrifuged at 12,000 x g at 4°C. Histones were washed one time in ice-cold 0.3 M HCl in acetone and two times ice-cold in 100% acetone before drying and resuspended in 20 mM Tris-HCl (pH 8.0), 0.4 N NaOH, supplemented with protease inhibitors [7.5 μM aprotinin, 0.5 mM leupeptin, 250 μM bestatin, 25 mM AEBSF-HCl]. Protein lysate concentration was determined by Bradford Assay (BioRad). 25 µg of acid-extracted proteins prepared in 1X LDS sample buffer (Invitrogen) containing 10% beta-mercaptoethanol were separated using SDS-PAGE. Proteins were transferred to nitrocellulose membranes and blocked in TBST + 5% milk. After primary and secondary antibody incubation, proteins of interest were visualized and quantified (Odyssey, Li-Cor).

For human and yeast studies, the primary antibodies used were as follows: H4K5ac (CST #8647; 1:1000), H4K8ac (CST #2594; 1:1000), H4K12ac (CST #13944; 1:1000), H4K16ac (Abcam #ab109463; 1:1000), H4K5/8/12/16ac (Abcam #ab177790; 1:1000), H4 (Cell Signaling Technology #2935; 1:1000), H3 (Abcam 1791; 0.1 µg/ml), TY1 (Diagenode C15200054; 2.2 µg/ml), HA (CST 6E2 1:1000). Secondary antibodies include the fluorophore-conjugated goat anti-rabbit 680 IgG (LiCor, 925-68071; 1:10,000) and goat anti-mouse 800 IgG (LiCor, 926-32210; 1:10,000).

### Quantitation of histone immunoblotting

The protein band intensities from immunoblots were quantitated using ImageJ software (Fiji), and mean fold changes in protein levels were calculated in Microsoft Excel (Microsoft Corporation). The protein band intensity was normalized for each endogenous histone PTM level to the total endogenous H4 band. The mean fold change was further calculated related to the wildtype H4 sample for each PTM. The data were graphed in GraphPad Prism 11 (GraphPad Software, LLC) as an average fold change compared to wildtype with standard deviation (n=3).

### Clonogenic assay

To measure cell clonogenicity, 500 HMEC cells/well were seeded in a 6-well plate in technical duplicate and incubated for a total of 21 days with media change every 3-4 days, after which cells were fixed in 50% EtOH and 10% acetic acid and crystal violet stained in 0.2% w/v crystal violet in 10% EtOH. Plates were imaged, destained in destaining solution (40% EtOH and 10% acetic acid), and crystal violet quantitated at OD_595_ (Biotek). Results were graphed in GraphPad Prism 11 (GraphPad Software, LLC), and statistical significance was assessed via one-way ANOVA analyses.

## Results

### Eaf1 overexpression rescues growth defects in H3K36 mutant cells

A previous high copy suppressor screen identified Esa1, the catalytic subunit of the NuA4 lysine acetyltransferase complex (Fig. 1A) (Clarke et al. 1999), as a suppressor of growth defects in H3K36-mutant cells (Lemon et al. 2022). To ensure balanced levels of H3 and H4 histones, we engineered new yeast strains with H3K36R or the H3K36M oncohistone mutant integrated into the endogenous *HHT2* locus with the (*hht1-hhf1)Δ* locus deleted, so that the only H3 protein expressed was either H3K36R or H3K36M and assessed caffeine sensitivity (Fig. 1B). Consistent with previous studies (Lemon et al. 2022; McDaniel et al. 2017), H3K36R and H3K36M cells show slow growth on plates that contain caffeine (Fig. 1B). H3K36R and H3K36M both eliminate methylation at H3K36, but the H3K36R mutant retains the original positive charge at this residue. By assaying both mutants, the impact of lost methylation and charge disruption can be distinguished. The major histone mutant genotypes of yeast strains utilized in these studies are detailed in Table 1.

**Table 1.**

| Label | Genotype |
| --- | --- |
| H3K36R | *hht2-K36R; (hht1-hhf1)Δ* |
| H3K36M | *hht2-K36M; (hht1-hhf1)Δ* |
| H4-4K→A | *hhf2-K5A/K8A/K12A/K16A* |
| H4-4K→R | *hhf2-K5R/K8R/K12R/K16R* |
| H3 + H4 | *HHT2; HHF2* |
| *hhΔ* | *(hht1-hhf1)Δ* |

To define which NuA4 subunit(s) are necessary to confer this suppression, we considered the functional activity of the NuA4 complex subunits, which are depicted in Figure 1A. Eaf1 is the scaffold that connects the NuA4 submodules to one another (Auger et al. 2008), and Yng2 stimulates the catalytic activity of Esa1 (Boudreault et al. 2003; Selleck et al. 2005). Eaf3 is part of the NuA4 complex, but it is also present in the Rpd3 deacetylase complex (Gavin et al. 2002; Ho et al. 2002). As a control, we employed Gcn5, which is the catalytic subunit of a different lysine acetyltransferase complex, SAGA (Morris et al. 2007; Espinola-Lopez and Tan 2021). The genes encoding each of these subunits were cloned into high copy vectors and expressed in WT, H3K36R, or H3K36M cells, then cell growth in the presence of caffeine was assessed by serial dilution assay (Fig. 1C). We saw no change in growth in WT cells, demonstrating that in no case does overexpression of these proteins non-specifically enhance rates of cell growth. As reported previously (Lemon et al. 2022), Esa1 improves growth in both H3K36R and H3K36M cells. Notably, the scaffold protein Eaf1 suppresses growth defects to an even greater extent than Esa1 in both H3 mutants. Eaf3, Yng2, and Gcn5 also suppress growth defects at a level similar to Esa1. Due to the strong growth of Eaf1-expressing cells, we focused our investigation on defining the mechanism of Eaf1-mediated suppression of H3K36 slow growth.

### The HSA domain in Eaf1 is necessary for rescue

To define how Eaf1 rescues cell growth, we created Eaf1 variants lacking specific domains (Fig. 2A) (Lu et al. 2022). The Eaf1 protein contains both an HSA domain and a SANT domain. HSA stands for helicase-SANT-associated domain, a domain that was originally found in several chromatin-regulatory proteins, often near SANT-related domains (Szerlong et al. 2008). The HSA domain of Eaf1 links Eaf1 to picNuA4 via interactions with Epl1 (Boudreault et al. 2003; Auger et al. 2008; Wang et al. 2018). Deleting the HSA domain disrupts complex stability and inhibits acetylation of lysine residues in H4 and H2A.Z (Lu et al. 2022). The Eaf1 SANT domain, named for Swi3, Ada2, N-CoR, and TFIIIB, participates in interaction with Tra1 as part of the Eaf1 scaffold function within NuA4 (Zukin et al. 2022). When the SANT domain is deleted from Eaf1, complex integrity and H4 and H2A.Z acetylation are not impacted (Wang et al. 2018; Lu et al. 2022). To investigate these Eaf1 variants, *EAF1* was deleted in the H3K36R, H3K36M, and control backgrounds. Cells lacking Eaf1 show slow growth (Auger et al. 2008) and sensitivity to caffeine (Fig. 2B). A lower concentration of caffeine was used for these experiments compared to other experiments (Fig. 1) because of the caffeine sensitivity of *eaf1*Δ mutants. The slow growth and caffeine sensitivity of H3K36R and H3K36M cells are exacerbated when *EAF1* is also deleted, as observed both in control rich media (Fig. 2B) and selective media (Fig. S2). Low copy plasmids expressing the Eaf1 domain variants were transformed into the *eaf1*Δ cells and cell growth was examined via serial dilution growth assay. The slow growth and caffeine sensitive phenotypes were rescued in the samples expressing Eaf1 or eaf1-SANTΔ, where the HSA domain is intact (Fig. 2C). Interestingly, eaf1-HSAΔ and eaf1-HSAΔSANTΔ confer slightly worse growth in the H3K36R and H3K36M cells compared to the vector only cells. The presence or absence of the SANT domain had no impact on cell growth. Based on this result, we conclude that the Eaf1 HSA domain is necessary to suppress the caffeine sensitive slow growth of H3K36 mutant yeast cells. We also tested whether Esa1 overexpression rescues growth defects in *eaf1*Δ cells, but we did not detect any rescue of the slow growth phenotype (Fig. 2C).

**Figure 2.**
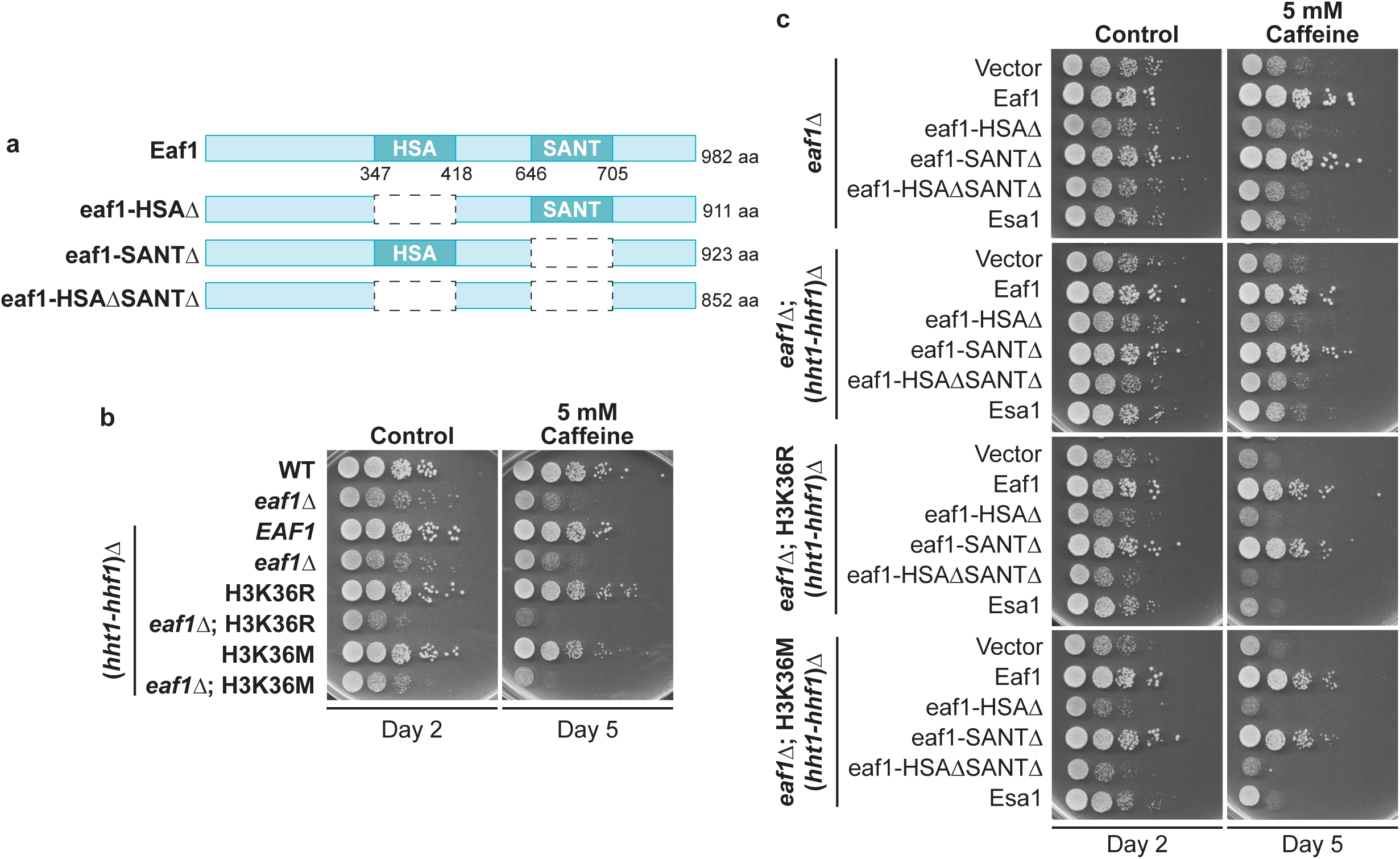
Loss of Eaf1 exacerbates growth defects on caffeine in H3K36 mutant cells, and the Eaf1 HSA domain is required to rescue H3K36 mutant growth defects. a) The Eaf1 protein contains an HSA domain and a SANT domain. A diagram of the Eaf1 protein domain variants utilized in this analysis is shown. b) Deleting *EAF1* in each histone H3 background causes a slow growth phenotype and caffeine sensitive growth. c) The Eaf1 variants depicted were expressed on low copy vectors in the *eaf1Δ* cells. Only the variants containing the HSA domain support growth on caffeine. Alt text: a) A diagram displaying the Eaf1 mutants with deleted domains as empty boxes. b) Yeast growth assays displaying that *eaf1*Δ cells of varying genetic backgrounds have slow growth and caffeine sensitivity. c) Yeast growth assays showing that the only cells that grow express Eaf1 containing the HSA domain.

### Histone H4 tail mutant cells show impaired viability

Given that the NuA4 complex is well characterized for its role in acetylating four lysines (H4K5, H4K8, H4K12, and H4K16) on the histone H4 N-terminal tail (Downey et al. 2015; Allard et al. 1999), mutants were created to eliminate acetylation at all four residues. Together, H4K5/8/12/16 were altered to either alanine or arginine. While neither alanine nor arginine can be acetylated, substitution of arginine at these residues maintains the original positive charge, providing the potential to distinguish between the absence of acetylation and change in residue charge. Budding yeast has two genes encoding histone H4, *HHF1* and *HHF2*. The H4-4K→A mutant represents the genotype *hhf2*-K5A/K8A/K12A/K16A, and the H4-4K→R mutant represents the genotype *hhf2*-K5R/K8R/K12R/K16R (Table 1). Due to conflicting prior reports on whether these H4 mutants can support viability when expressed as the only H4 copy (Megee et al. 1990; Glowczewski et al. 2004), we tested whether the H4-4K→A and H4-4K→R mutants were viable alone or in combination with H3K36R or H3K36M (Fig. 3B). Similar to prior studies, endogenous (*hht1-hhf1*)Δ was deleted for this analysis. The H3 mutant is incorporated at the *hht2* gene, and the H4 mutant is incorporated at the *hhf2* gene. The cells have a WT copy of H3 (*HHT2*) and H4 (*HHF2*) expressed on a low copy plasmid with a *URA3* marker. When grown on selective media lacking uracil (Ura-), cells retain this plasmid and can grow. However, when cells are grown on media containing 5-fluoroorotic acid (5-FOA), they must lose the *URA3*-containing vector to survive (Boeke et al. 1987). Therefore, colonies that grow on 5-FOA plates are viable without exogenous WT H3 and H4 (Fig. 3A). The number of colonies that grew on either Ura- or 5-FOA plates were counted, and the ratio of colonies on 5-FOA plates to colonies on Ura- plates for each genotype is shown in Figure 3B. Representative images of plates are presented in Figure S3. Control cells that express H4-4K→A and contain endogenous WT *HHT1-HHF1* showed the highest level of viability, with a mean ratio of 0.254. When (*hht1-hhf1*)Δ was deleted, so that H4-4K→A is the only histone H4 expressed, the ratio reduces significantly to 0.031 (p<0.0001). When combined with H3K36R, viability was further significantly reduced (p<0.0001), and when combined with H3K36M, no colonies survived (p<0.0001). The H4-4K→R mutant was even more detrimental than H4-4K→A when expressed as the only H4 histone (Fig. 3B). A few colonies survived on 5-FOA when H4-4K→R was combined with (*hht1-hhf1*)Δ alone or with H3K36M. However, some of these colonies were sequenced and found to have reverted both H3 (*HHT2*) and H4 (*HHF2*) to a WT sequence, suggesting that the actual viability may be lower. The growth of these cells was also investigated by serial dilution assay on Ura- and 5-FOA plates (Fig. 3C). The extent of growth when the H4-4K→A or H4-4K→R mutant is expressed in this experiment corresponds to what was observed in Fig. 3B. To confirm that H4-4K→A expression results in an absence of acetylation at the four N-terminal tail lysine residues, an immunoblot was performed on acid extracted histones from H4-4K→A cells that did not contain the plasmid expressing WT *HHT2-HHF2*. As expected in the H4-4K→A; (*hht1-hhf1*)Δ mutant, which lacks WT H4, acetylation at these residues is not detectable (Fig. 3D). A corresponding immunoblot with H4-4K→R; (*hht1-hhf1*)Δ cells could not be performed because no colonies lacking the *HHT2-HHF2* plasmid were identified that had not reverted the mutants back to WT sequence, so no viable yeast cells were available to analyze.

**Figure 3.**
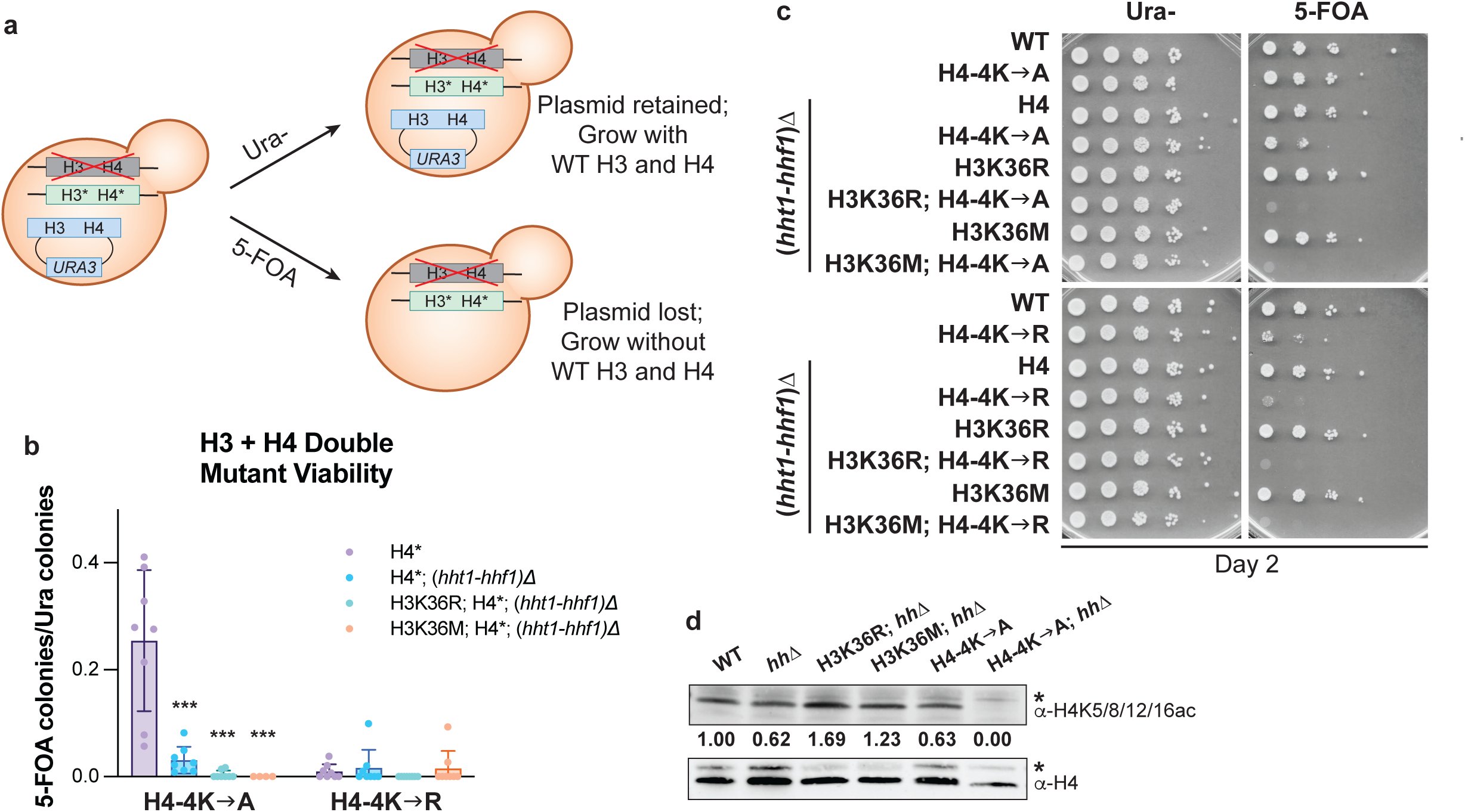
Altering histone H4 tail lysines to alanine or arginine severely impairs cell growth. a) A diagram illustrating the experimental strategy pursued is shown. Cells express mutant H3 from the endogenous *hht2* locus and mutant H4 from the endogenous *hhf2* locus, as denoted with *. The *hht1-hhf1* genes are deleted as in previous experiments. Cells contain an exogenous low copy *URA3* vector encoding wildtype *HHT2-HHF2*. When plated on URA- dropout media, all cells retain the *HHT2-HHF2* vector and grow normally. When plated on 5-FOA media, cells must lose the *URA3*-containing *HHT2-HHF2* vector to grow. Therefore, any surviving colonies are viable with the described mutant histones H3 and H4. b) For this and subsequent results, H4-4K→A represents the mutant *hhf2*-K5A/K8A/K12A/K16A, and H4-4K→R represents the mutant *hhf2*-K5R/K8R/K12R/K16R. Colonies were counted, and the ratio of double mutants surviving on 5-FOA to those on URA- is presented on the graph. *** represents p<0.0001. c) A serial dilution growth assay was employed to assess the growth of the H3/H4 double mutant cells on both Ura-and 5-FOA plates. d) A representative immunoblot analyzing H4K5/8/12/16ac signal is shown. The numbers below the blot represent the quantified H4K5/8/12/16ac band intensities for each sample normalized to their respective H4 band and relative to WT, set to 1.0. *Denotes non-specific bands for each antibody. Due to space constraints, *hh*Δ is used to denote (*hht1-hhf1*)Δ. Alt text: a) A diagram representing the experiment performed. A budding yeast cell expresses endogenous mutant histones H3 and H4 and exogenous plasmid with WT histones H3 and H4. The cell will retain the plasmid and grow when plated on Ura- media, and the cell will lose the plasmid when plated on 5-FOA but only survive if it is viable with the two histone mutants. b) Graphical representation of the double viability experiment showing that the H4-4K→A cells have the highest viability. c) Yeast growth assay showing that cell growth by serial dilution is similar to that of colony counts. d) Presence of the PTM H4K5/8/12/16ac is assessed by band intensity to show that the H4-4K→A mutant lacks acetylation.

### Histone H4 mutants perturb acetylation and cause impaired growth even when WT histone H3 and H4 are expressed

As cells expressing histone H4 mutants show low viability (Fig. 3A-C), a low copy vector expressing WT H3 and H4 (*HHT2-HHF2*) was transformed into control and mutant strains, similar to the experimental set up depicted in Figure 3A. Despite the presence of a WT copy of each H3 and H4, both H4-4K→A and H4-4K→R mutants still exhibit slow growth on control plates and sensitivity to caffeine (Fig. 4A). These phenotypes reveal that the H4 mutants act in a dominant manner. Furthermore, when combined with the H3K36M mutant, both H4-4K→A and H4-4K→R mutants display an additive impact on caffeine sensitivity even in the presence of WT H3 and H4 (Fig. 4A).

**Figure 4.**
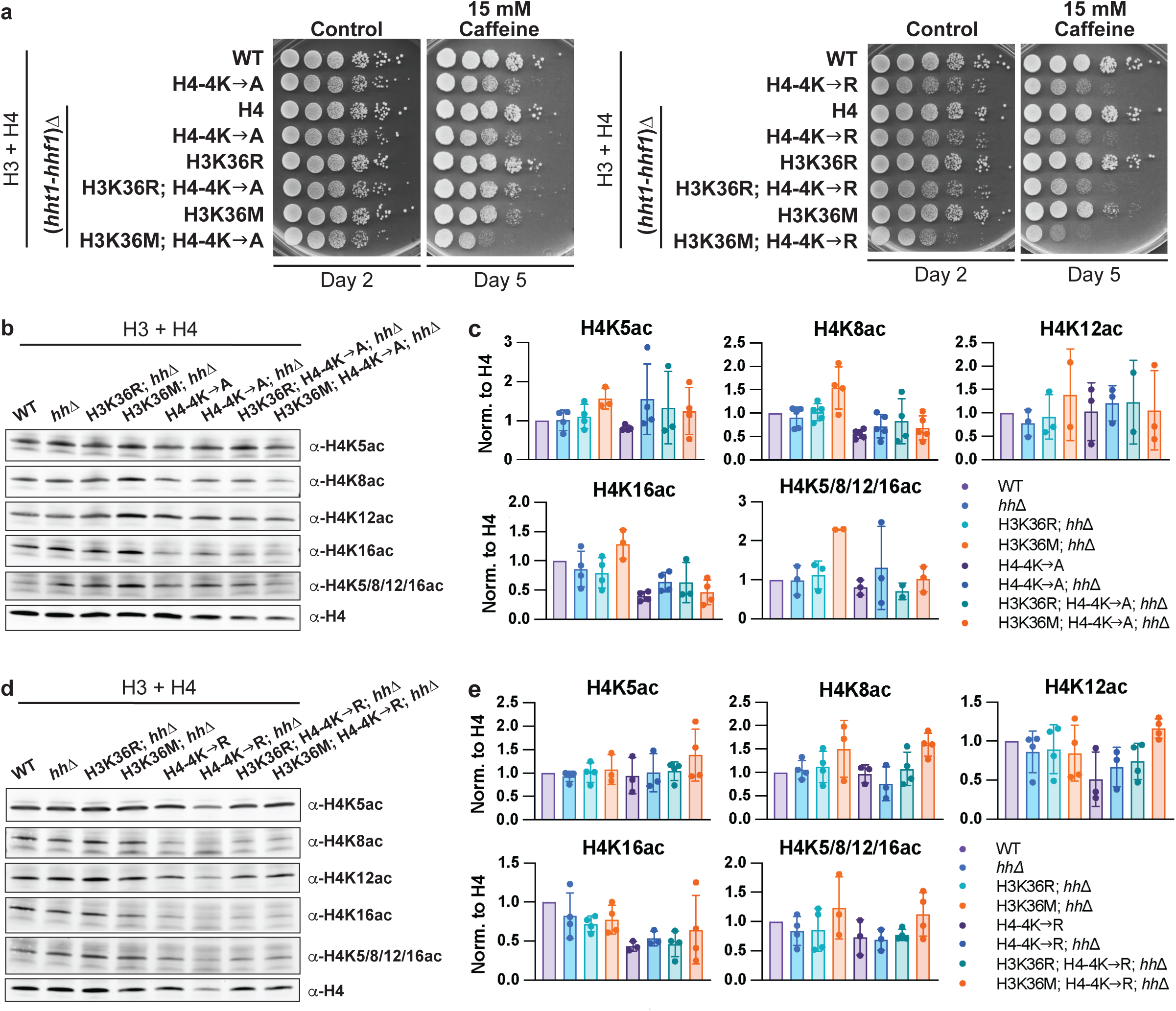
When WT H3 and H4 are present, H4 tail acetylation trends lower in H4-4K→A and H4-4K→R cells, and these H4 mutants also display growth defects. a) Serial dilution growth assays reveal that both H4-4K→A and H4-4K→R mutants have a slow growth phenotype and sensitivity to caffeine. b) Given the severe growth defect in double H3 and H4 mutants, the following experiments were conducted in the presence of an exogenous wildtype H3 and H4 (*HHT2-HHF2*) low copy vector, represented by the label “H3 + H4”. A representative immunoblot of the H4-4K→A mutant cells, which was probed to detect the indicated lysine acetylation marks, is shown. Due to space constraints, *hh*Δ is used to denote (*hht1-hhf1*)Δ. c) Quantification of the immunoblots in panel B, with each mark normalized to H4, which was set to 1.0. Graphs display the fold change compared to each respective WT. d) A representative immunoblot of the H4-4K→R mutant cells. e) Quantification of the immunoblots shown in panel D, with each mark normalized to H4, which was set to 1.0. Graphs display the fold change compared to the respective WT. Alt text: a) Yeast growth assays showing that the H4-4K→A and H4-4K→R cells with varying genetic backgrounds have slow growth and caffeine sensitivity. b) Assessment of various histone H4 acetylation events in H4-4K→A mutant cells by intensity of band signal. c) Graphs displaying the quantification of H4 acetylation in H4-4K→A cells. d) Assessment of various histone H4 acetylation events in H4-4K→R mutant cells by intensity of band signal. e) Graphs displaying the quantification of H4 acetylation in H4-4K→R cells.

Next, to characterize the impact of combined histone H3 and H4 mutants on global H4 acetylation, immunoblots of acid extracted histone lysates were performed. Acetylation of the lysines that are the histone H4 targets of NuA4 was examined using antibodies that recognize the four acetyl marks [H4K(5/8/12/16)ac] individually or in combination (Fig. 4B). For cells expressing the H4-4K→A mutant, there was variability in the impact on H4 acetylation (Fig. 4C). Comparing the H4-4K→A signal to that of the WT cells, acetylation trended lower when samples were probed to detect H4K5ac, K8ac, K16ac, or H4K(5/8/12/16)ac. This finding suggests that the H4-4K→A mutant alone reduces H4 acetylation at multiple lysine residues even when residual WT H4 is present. The H4-4K→A mutants in all four genetic backgrounds trend towards lower H4K8- and H4K16-acetylation (Fig. 4C). For cells expressing H4-4K→R, a similar decrease in acetylation in the H4-4K→R mutant alone compared to WT was observed at H4K12 and H4K16, though acetylation was similar to WT at other residues (Fig. 4D, E). A trend towards reduced acetylation in all four H4-4K→R genetic backgrounds was also observed for H4K16. Interestingly, H4K16 is typically the first residue to be acetylated in H4 tails for both yeast and mammals (Clarke et al. 1993; Munks et al. 1991), and the acetylation state of H4K16 functions as a switch between heterochromatin and euchromatin (Millar et al. 2004). Perturbations to H4K16ac may have severe consequences on epigenetic regulation.

### Growth defects in histone H4 mutants are not rescued by Esa1 or Eaf1

As methylation at H3K36 assists in NuA4 nucleosomal recruitment (Ginsburg et al. 2014), the absence of H3K36 methylation in H3K36R and H3K36M mutant cells may be rescued because overexpression of NuA4 subunits compensates for the loss of H4 N-terminal tail acetylation. To test this hypothesis, we examined whether overexpression of Esa1 or Eaf1 could suppress growth defects in the H4-4K→A or H4-4K→R mutant cells (Fig. 5, S5). As shown in Fig. 1C, Esa1 and Eaf1 rescue growth defects in H3K36R or H3K36M mutant cells (Fig. 5). However, when H4-4K→A or H4-4K→R is combined with either H3K36R or H3K36M, neither Esa1 nor Eaf1 can rescue growth defects (Fig. 5). This result is consistent with the hypothesis that suppression of growth defects in H3K36M and H3K36R cells by NuA4 components is dependent on acetylation at histone H4 tails.

**Figure 5.**
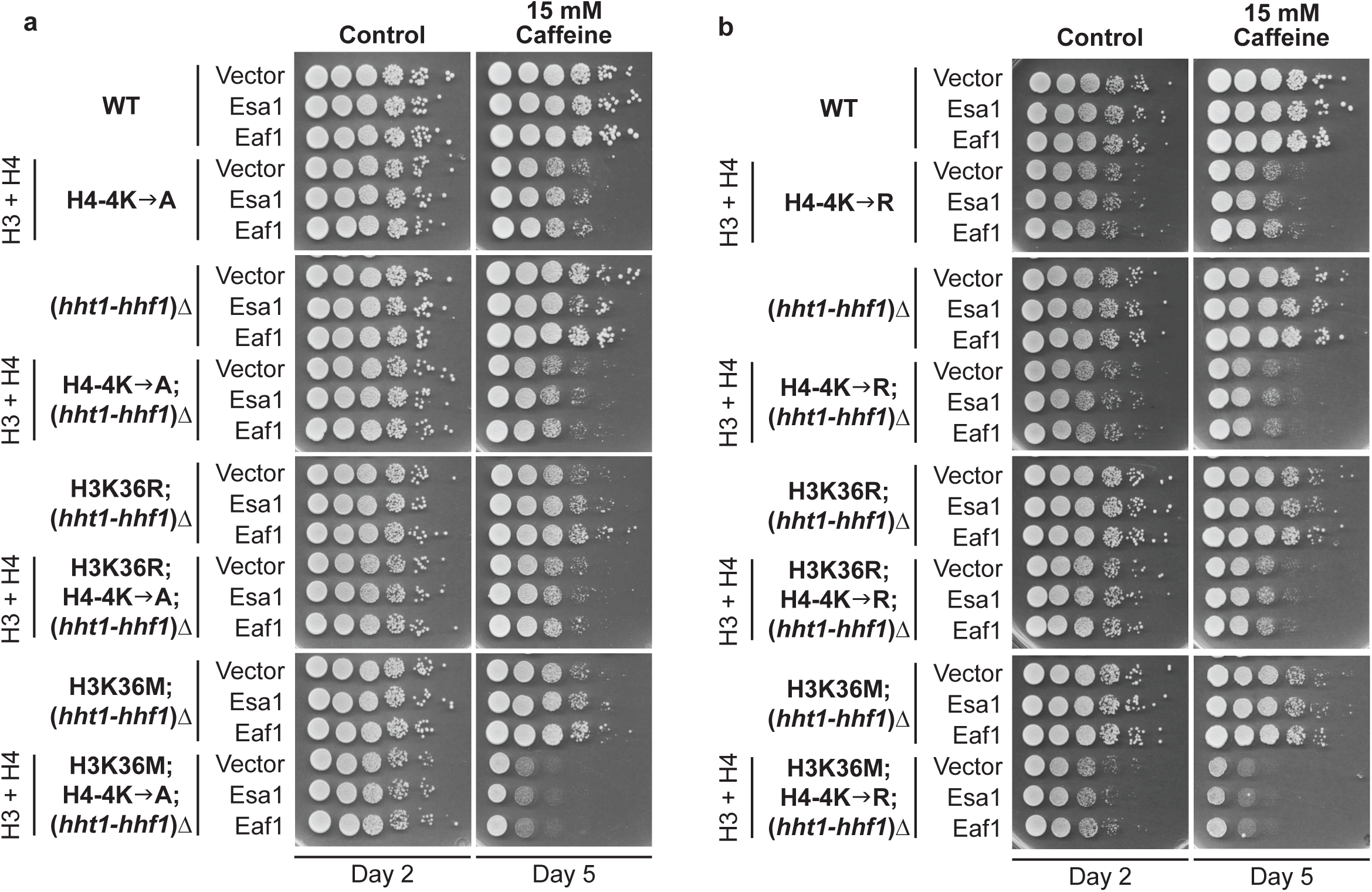
Histone H4 tail lysines are required for Esa1 and Eaf1 to suppress growth defects in H3K36R/M cells. a) H4-4K→A cells were transformed with high copy Esa1- or Eaf1-plasmids or the Vector control. A serial dilution growth assay was used to assess the Esa1- or Eaf1-meduated suppression of H3K36M mutant cells in combination with the H4 acetylation mutants. b) A serial dilution growth assay was used to assess the Esa1- or Eaf1-meduated suppression of H3K36R mutant cells in combination with the H4 acetylation mutants. Alt text: a) Yeast growth assays showing that cells expressing H4-4K→A lack improved growth after Esa1 or Eaf1 overexpression. b) Yeast growth assays showing that cells expressing H4-4K→R lack improved growth after Esa1 or Eaf1 overexpression.

### Altering NuA4 function in human H3K36M cells impairs clonogenic potential

As these investigations were performed with the H3K36M oncohistone mutant, which is associated with multiple cancer types in humans (Behjati et al. 2013; Papillon-Cavanagh et al. 2017), we tested whether we could translate our findings in budding yeast to a human system. We previously established immortalized but untransformed human mammary epithelial cell (HMEC) lines transduced with C-terminal TY1-tagged WT H3 or H3K36M (Sad et al. 2025) (Fig. 6A). This oncohistone model can be used to assess how certain perturbations influence growth of H3K36M cells. The human homolog of Esa1 is the histone acetyltransferase TIP60 (Doyon et al. 2004), and the small molecule inhibitor MG149 selectively targets the MYST family of histone acetyltransferases including TIP60 and MOF (Ghizzoni et al. 2012). TIP60 inhibition via MG149 exhibits preclinical anticancer activity in models of anaplastic thyroid cancer and mesothelioma (Cregan et al. 2016; Ning et al. 2022; Sheng et al. 2025). In yeast cells, overexpression of NuA4 components improves cell growth of the H3K36M and H3K36R models (Fig. 1C). In HMEC cells expressing WT H3 or the oncohistone H3K36M, we tested whether TIP60 inhibition selectively impairs H3K36M cell growth. We employed a limited dilution, also known as a clonogenic assay, which assesses the transformed phenotype of cells growing into isolated colonies after being seeded at low density (Fig. 6B). Neither 0.5 μM or 1 μM MG149 impact the clonogenicity of WT H3 cells (Fig. 6C). However, TIP60 inhibition (1 μM) significantly reduced clonogenic growth of H3K36M cells (from 0.978 +/- 0.26 to 0.56 +/- 0.11; *p* = 0.02) (Fig. 6C). These data suggest that inhibition of the NuA4 human homologue and histone acetyltransferase TIP60 specifically reduces growth in cells expressing the oncohistone H3K36M.

**Figure 6.**
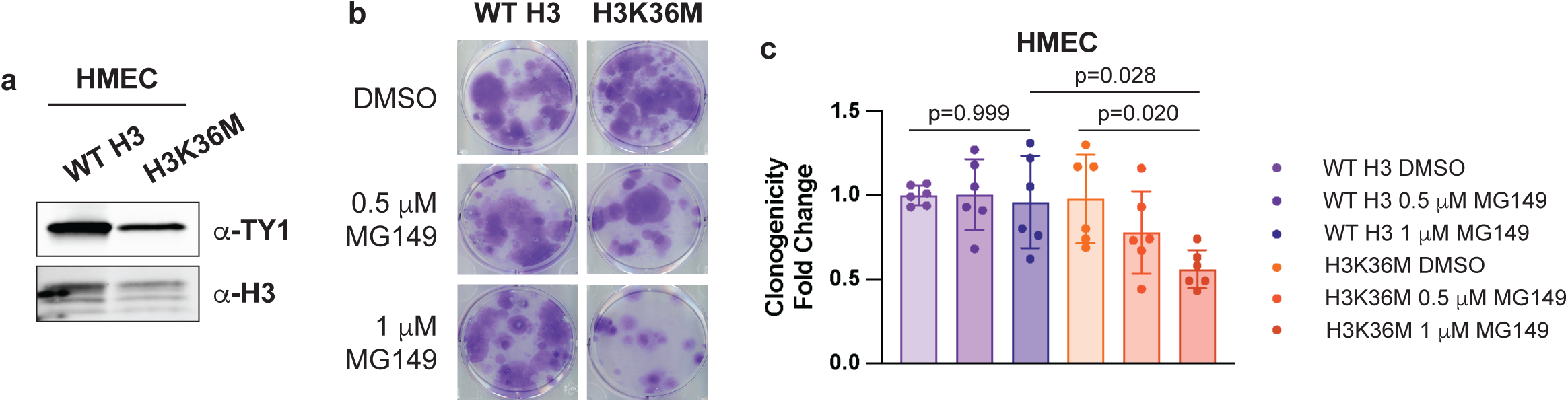
H3K36M-expressing human cells have decreased clonogenicity when NuA4 activity is inhibited. a) Immunoblot displaying the ectopic expression of TY1-tagged WT H3 and H3K36M in an immortalized, non-transformed human breast cell line (HMEC). b) Representative wells from the clonogenic assay. Cells were seeded at low density and grown for 21 days in DMSO (vehicle) or varying concentrations of MG149 (TIP60 inhibitor). Then, cells were fixed and stained with crystal violet. c) Quantification of crystal violet staining from panel B. Absorbance was normalized to the respective DMSO sample for each cell line, and the graph displays fold change. A one-way ANOVA was performed to assess significance. Alt text: a) Bands of TY1-tagged exogenous WT H3 and H3K36M. b) Wells from a clonogenic assay where the clonal cell populations are stained purple with crystal violet. c) A graph representing quantification of the staining in the clonogenic assay represented as fold change.

## Discussion

In this work, we characterize Eaf1 as a novel suppressor of caffeine-sensitive growth of H3K36M and H3K36R yeast cells. We demonstrate that this suppression is dependent on lysine acetylation within H4. We extend our analysis to human cells and demonstrate that inhibiting the NuA4/TIP60 acetyltransferase preferentially decreases the clonogenic potential of an H3K36M oncohistone model as compared to control cells. Taken together, these studies provide insight into how oncohistones may alter cell growth and provide a potential new therapeutic target for oncohistone-driven cancers.

We initially identified Esa1 as a suppressor of growth defects in these same H3K36 mutants (Lemon et al. 2022), and we were surprised to find that Eaf1 confers even stronger rescue than Esa1 (Fig. 1C). We also discovered that this rescue of impaired growth is dependent on the presence of the HSA domain in Eaf1 (Fig. 2C). The Eaf1 HSA domain is necessary for maintaining NuA4 complex integrity through its interactions with other NuA4 subunits (Lu et al. 2022). While previous work shows that eaf1-HSAΔ mutants lack an intact complete NuA4 complex, they do retain all members of the picNuA4 complex (Lu et al. 2022). Therefore, the complete NuA4 complex may be required for suppression of H3K36R and H3K36M growth defects, suggesting that picNuA4 is not sufficient to rescue growth defects of H3K36 mutant cells. This hypothesis may also explain our observation that Eaf1 exhibits stronger growth suppression than Esa1 (Fig. 1C) as Eaf1 is specific to the complete NuA4 complex, while Esa1 is present in both NuA4 and picNuA4. Thus, Eaf1 overexpression may increase the proportion or bias the formation of the complete NuA4 complex compared to picNuA4.

We also characterized H4-4K→A and H4-4K→R mutants alone or in combination with H3K36R and H3K36M. When the H4-4K→A mutant is present with at least one copy of a WT histone H4 gene, yeast cells are viable (Fig. 3B, C). However, deleting the second histone H4 gene encoded in the *S. cerevisiae* genome severely impairs viability in this context. Adding the H3K36R or H3K36M mutant to this background further reduces viability, revealing that there is an additive effect when the histone H3K36 oncohistone mutants and H4 acetylation impaired mutants are combined. There were no surviving colonies with the H3K36M + H4-4K→A double mutants. The only four H3K36R + H4-4K→A double mutant colonies that were identified in the viability assay (Fig. 3B) were Sanger sequenced at the H3 and H4 genes, and these colonies maintained mutant H3 and H4 sequences (data not shown). However, we cannot rule out the possibility that these colonies acquired a background mutation that allowed growth despite the double histone H3 + H4 mutant cells. The H4-4K→R mutant severely impaired viability in all genetic backgrounds (Fig. 3B, C). Sequencing the H3 and H4 genes in a subset of the surviving colonies revealed that the H4-4K→R cells, which express WT H4 at the second endogenous copy (*HHF1*), retained the H4-4K→R mutant. However, the H4-4K→R; (*hht1-hhf1*)Δ cells reverted to WT H4 (*HHF2*). Interestingly, all surviving colonies in double H3 + H4 mutant backgrounds had reverted to WT sequences at both H3 (*HHT2*) and H4 (*HHF2*) genes. Because there were few H4-4K→R colonies on 5-FOA, and because the double mutant colonies had acquired revertant mutations, it is likely that the H4-4K→R mutant has a stronger impact on viability than the H4-4K→A does. Interestingly, despite being more detrimental to viability, the arginine mutant retains the original amino acid charge. It is possible, therefore, that the H4-4K→R mutant binds histone modifying enzymes more tightly than the H4-4K→A mutant does, effectively titrating these key enzymes and broadly impairing their function.

Given their impaired viability, we provided a vector with WT H3 (*HHT2*) and H4 (*HHF2*) in all H4-4K→A and H4-4K→R mutants for subsequent experiments. We found that despite the presence of WT histone H3 and H4, cells with H4-4K→A or H4-4K→R alone or in combination with H3K36R or H3K36M showed slow growth and caffeine sensitivity (Fig. 4E). These data reveal that both the H4-4K→A and H4-4K→R mutants are dominant to WT H4. Oncohistone mutants including H3K36M and H3K27M drive cancer in a dominant negative manner because cancer occurs even though only a single copy of multiple histone genes is mutated. Prior studies have shown that H3K36M and H3K27M bind methyltransferases that catalyze H3K36 and H3K27 methylation, SETD2 and EZH2, respectively, more tightly than WT H3 binds to the same respective enzyme, altering the epigenetic landscape due to impaired modification of H3K36 and/or H3K27 (Fang et al. 2016; Lu et al. 2016; Justin et al. 2016). The trend towards decreased H4 acetylation observed in Fig. 4B and D may be consistent with the proportion of WT to mutant H4 in each genotype. However, a dominant negative interaction may cause the decrease in H4 acetylation. Further work is required to characterize the mechanism of dominance in the H4-4K→A and H4-4K→R cells.

A critical finding for this investigation is that Eaf1 and Esa1 do not suppress growth defects in cells expressing the H4-4K→A or H4-4K→R mutant (Fig. 5). This result is consistent with the hypothesis that NuA4 suppression of mutant H3K36 growth defects is dependent on NuA4-mediated acetylation at histone H4 tails. Interestingly, all mutant H4 cells tested expressed a plasmid with a WT H3 and H4 gene, meaning that H4 proteins with the capacity to be acetylated were present in these cells. Despite expressing WT H4, Eaf1 and Esa1 were still unable to rescue growth defects in H4 mutant cells. Given the immunoblot results in Fig. 4, the histone H4 acetylation event that is most consistently decreased in the H4 mutant cells is H4K16ac; therefore, rescue by Eaf1 or Esa1 may be dependent specifically on the H4K16ac mark.

Finally, we translated our findings to a human experimental system. We found that inhibition of the NuA4/TIP60 complex reduces clonogenic potential in human cells that express the H3K36M oncohistone (Fig. 6). This finding highlights the power of model systems such as *S. cerevisiae* that are amenable to rapid discovery through genetic screens to provide a platform to guide future studies. This finding is also in line with our hypothesis that increasing acetylation at histone H4 N-terminal tails rescues growth defects caused by H3K36M. When NuA4 activity is decreased in H3K36M human cells via TIP60 inhibition, cellular growth is impaired. Therefore, TIP60 inhibition may be a potential avenue for therapeutic exploration in cancers expressing the H3K36M oncohistone.

### Data Availability Statement

The data underlying this article will be shared on reasonable request to the corresponding author.

## Acknowledgments

We would also like to thank Dr. Michael Kobor and his lab manager for providing us with the Eaf1 domain deletion plasmids (Lu et al. 2022). Finally, we would like to thank members of the Spangle and Corbett lab as well as our Team Histone undergraduates for providing support and feedback on specific experiments.

## Author Contributions

CYJ conceptualized the project, designed and performed experiments, and wrote the manuscript. LC performed experiments, and MF created crucial reagents. JMS and AHC provided guidance over experimental design and interpretation and edited the manuscript.

## References

1. Adams, A, DE Gottschling, and CA Kaiser. Stearns. T. (1997) Methods in Yeast Genetics. Cold Spring Harbor Laboratory Press, Cold Spring Harbor, NY.

2. Allard, S., R. T. Utley, J. Savard et al. 1999. “Nua4, an Essential Transcription Adaptor/Histone H4 Acetyltransferase Complex Containing Esa1p and the Atm-Related Cofactor Tra1p.” Embo j 18 (18):5108–19. 10.1093/emboj/18.18.5108.

3. Auger, A., L. Galarneau, M. Altaf et al. 2008. “Eaf1 Is the Platform for Nua4 Molecular Assembly That Evolutionarily Links Chromatin Acetylation to Atp-Dependent Exchange of Histone H2a Variants.” Mol Cell Biol 28 (7):2257–70. 10.1128/mcb.01755-07.

4. Behjati, S., P. S. Tarpey, N. Presneau et al. 2013. “Distinct H3f3a and H3f3b Driver Mutations Define Chondroblastoma and Giant Cell Tumor of Bone.” Nat Genet 45 (12):1479–82. 10.1038/ng.2814.

5. Boeke, J. D., J. Trueheart, G. Natsoulis, and G. R. Fink. 1987. “5-Fluoroorotic Acid as a Selective Agent in Yeast Molecular Genetics.” Methods Enzymol 154:164–75. 10.1016/0076-6879(87)54076-9.

6. Boudreault, A. A., D. Cronier, W. Selleck et al. 2003. “Yeast Enhancer of Polycomb Defines Global Esa1-Dependent Acetylation of Chromatin.” Genes Dev 17 (11):1415–28. 10.1101/gad.1056603.

7. Clarke, A. S., J. E. Lowell, S. J. Jacobson, and L. Pillus. 1999. “Esa1p Is an Essential Histone Acetyltransferase Required for Cell Cycle Progression.” Mol Cell Biol 19 (4):2515–26. 10.1128/mcb.19.4.2515.

8. Clarke, D. J., L. P. O’Neill, and B. M. Turner. 1993. “Selective Use of H4 Acetylation Sites in the Yeast Saccharomyces Cerevisiae.” Biochem J 294 ( Pt 2) (Pt 2):557–61. 10.1042/bj2940557.

9. Cregan, S., L. McDonagh, Y. Gao et al. 2016. “Kat5 (Tip60) Is a Potential Therapeutic Target in Malignant Pleural Mesothelioma.” Int J Oncol 48 (3):1290–6. 10.3892/ijo.2016.3335.

10. Downey, M., J. R. Johnson, N. E. Davey et al. 2015. “Acetylome Profiling Reveals Overlap in the Regulation of Diverse Processes by Sirtuins, Gcn5, and Esa1.” Mol Cell Proteomics 14 (1):162–76. 10.1074/mcp.M114.043141.

11. Doyon, Y., W. Selleck, W. S. Lane, S. Tan, and J. Côté. 2004. “Structural and Functional Conservation of the Nua4 Histone Acetyltransferase Complex from Yeast to Humans.” Mol Cell Biol 24 (5):1884–96. 10.1128/mcb.24.5.1884-1896.2004.

12. Duina, Andrea A, and Claire E Turkal. 2017. “Targeted in Situ Mutagenesis of Histone Genes in Budding Yeast.” Journal of Visualized Experiments: JoVE (119):55263.

13. Espinola-Lopez, J. M., and S. Tan. 2021. “The Ada2/Ada3/Gcn5/Sgf29 Histone Acetyltransferase Module.” Biochim Biophys Acta Gene Regul Mech 1864 (2):194629. 10.1016/j.bbagrm.2020.194629.

14. Fang, D., H. Gan, J. H. Lee et al. 2016. “The Histone H3.3k36m Mutation Reprograms the Epigenome of Chondroblastomas.” Science 352 (6291):1344–8. 10.1126/science.aae0065.

15. Friis, R. M., B. P. Wu, S. N. Reinke, D. J. Hockman, B. D. Sykes, and M. C. Schultz. 2009. “A Glycolytic Burst Drives Glucose Induction of Global Histone Acetylation by Picnua4 and Saga.” Nucleic Acids Res 37 (12):3969–80. 10.1093/nar/gkp270.

16. Gavin, Anne-Claude, Markus Bösche, Roland Krause, et al. 2002. “Functional Organization of the Yeast Proteome by Systematic Analysis of Protein Complexes.” Nature 415 (6868):141–47.

17. Ghizzoni, M., J. Wu, T. Gao, H. J. Haisma, F. J. Dekker, and Y. George Zheng. 2012. “6-Alkylsalicylates Are Selective Tip60 Inhibitors and Target the Acetyl-Coa Binding Site.” Eur J Med Chem 47 (1):337–44. 10.1016/j.ejmech.2011.11.001.

18. Ginsburg, D. S., T. E. Anlembom, J. Wang, S. R. Patel, B. Li, and A. G. Hinnebusch. 2014. “Nua4 Links Methylation of Histone H3 Lysines 4 and 36 to Acetylation of Histones H4 and H3.” J Biol Chem 289 (47):32656–70. 10.1074/jbc.M114.585588.

19. Ginsburg, D. S., C. K. Govind, and A. G. Hinnebusch. 2009. “Nua4 Lysine Acetyltransferase Esa1 Is Targeted to Coding Regions and Stimulates Transcription Elongation with Gcn5.” Mol Cell Biol 29 (24):6473–87. 10.1128/mcb.01033-09.

20. Glowczewski, L., J. H. Waterborg, and J. G. Berman. 2004. “Yeast Chromatin Assembly Complex 1 Protein Excludes Nonacetylatable Forms of Histone H4 from Chromatin and the Nucleus.” Mol Cell Biol 24 (23):10180–92. 10.1128/mcb.24.23.10180-10192.2004.

21. Hill, J. E., A. M. Myers, T. J. Koerner, and A. Tzagoloff. 1986. “Yeast/E. Coli Shuttle Vectors with Multiple Unique Restriction Sites.” Yeast 2 (3):163–7. 10.1002/yea.320020304.

22. Ho, Yuen, Albrecht Gruhler, Adrian Heilbut, et al. 2002. “Systematic Identification of Protein Complexes in Saccharomyces Cerevisiae by Mass Spectrometry.” Nature 415 (6868):180–83.

23. Jacquet, K., A. Fradet-Turcotte, N. Avvakumov et al. 2016. “The Tip60 Complex Regulates Bivalent Chromatin Recognition by 53bp1 through Direct H4k20me Binding and H2ak15 Acetylation.” Mol Cell 62 (3):409–21. 10.1016/j.molcel.2016.03.031.

24. Johnson, Paige, Virginia Mitchell, Kelsi McClure et al. 2015. “A Systematic Mutational Analysis of a Histone H3 Residue in Budding Yeast Provides Insights into Chromatin Dynamics.” G3: Genes, Genomes, Genetics 5 (5):741–49.

25. Jones, R. B., J. Farhi, M. Adams et al. 2022. “Targeting Mll Methyltransferases Enhances the Antitumor Effects of Pi3k Inhibition in Hormone Receptor-Positive Breast Cancer.” Cancer Res Commun 2 (12):1569–78. 10.1158/2767-9764.Crc-22-0158.

26. Justin, N., Y. Zhang, C. Tarricone et al. 2016. “Structural Basis of Oncogenic Histone H3k27m Inhibition of Human Polycomb Repressive Complex 2.” Nat Commun 7:11316. 10.1038/ncomms11316.

27. Lemon, L. D., S. Kannan, K. W. Mo et al. 2022. “A Saccharomyces Cerevisiae Model and Screen to Define the Functional Consequences of Oncogenic Histone Missense Mutations.” G3 (Bethesda) 12 (7) 10.1093/g3journal/jkac120.

28. Lewis, P. W., M. M. Müller, M. S. Koletsky et al. 2013. “Inhibition of Prc2 Activity by a Gain-of-Function H3 Mutation Found in Pediatric Glioblastoma.” Science 340 (6134):857–61. 10.1126/science.1232245.

29. Li, L., and Y. Wang. 2017. “Cross-Talk between the H3k36me3 and H4k16ac Histone Epigenetic Marks in DNA Double-Strand Break Repair.” J Biol Chem 292 (28):11951–59. 10.1074/jbc.M117.788224.

30. Lin, Y. Y., J. Y. Lu, J. Zhang et al. 2009. “Protein Acetylation Microarray Reveals That Nua4 Controls Key Metabolic Target Regulating Gluconeogenesis.” Cell 136 (6):1073–84. 10.1016/j.cell.2009.01.033.

31. Lu, C., S. U. Jain, D. Hoelper et al. 2016. “Histone H3k36 Mutations Promote Sarcomagenesis through Altered Histone Methylation Landscape.” Science 352 (6287):844–9. 10.1126/science.aac7272.

32. Lu, Phoebe YT, Alyssa C Kirlin, Maria J Aristizabal et al. 2022. “A Balancing Act: Interactions within Nua4/Tip60 Regulate Picnua4 Function in Saccharomyces Cerevisiae and Humans.” Genetics 222 (3):iyac136.

33. Luger, K., A. W. Mäder, R. K. Richmond, D. F. Sargent, and T. J. Richmond. 1997. “Crystal Structure of the Nucleosome Core Particle at 2.8 a Resolution.” Nature 389 (6648):251–60. 10.1038/38444.

34. McDaniel, S. L., A. J. Hepperla, J. Huang et al. 2017. “H3k36 Methylation Regulates Nutrient Stress Response in Saccharomyces Cerevisiae by Enforcing Transcriptional Fidelity.” Cell Rep 19 (11):2371–82. 10.1016/j.celrep.2017.05.057.

35. Megee, P. C., B. A. Morgan, B. A. Mittman, and M. M. Smith. 1990. “Genetic Analysis of Histone H4: Essential Role of Lysines Subject to Reversible Acetylation.” Science 247 (4944):841–5. 10.1126/science.2106160.

36. Millar, C. B., S. K. Kurdistani, and M. Grunstein. 2004. “Acetylation of Yeast Histone H4 Lysine 16: A Switch for Protein Interactions in Heterochromatin and Euchromatin.” Cold Spring Harb Symp Quant Biol 69:193–200. 10.1101/sqb.2004.69.193.

37. Mitchener, M. M., and T. W. Muir. 2022. “Oncohistones: Exposing the Nuances and Vulnerabilities of Epigenetic Regulation.” Mol Cell 82 (16):2925–38. 10.1016/j.molcel.2022.07.008.

38. Morris, S. A., B. Rao, B. A. Garcia et al. 2007. “Identification of Histone H3 Lysine 36 Acetylation as a Highly Conserved Histone Modification.” J Biol Chem 282 (10):7632–40. 10.1074/jbc.M607909200.

39. Munks, R. J., J. Moore, L. P. O’Neill, and B. M. Turner. 1991. “Histone H4 Acetylation in Drosophila. Frequency of Acetylation at Different Sites Defined by Immunolabelling with Site-Specific Antibodies.” FEBS Lett 284 (2):245–8. 10.1016/0014-5793(91)80695-y.

40. Ning, J., Q. Sun, Z. Su et al. 2022. “The Ck1δ/ɛ-Tip60 Axis Enhances Wnt/β-Catenin Signaling Via Regulating β-Catenin Acetylation in Colon Cancer.” Front Oncol 12:844477. 10.3389/fonc.2022.844477.

41. Papillon-Cavanagh, S., C. Lu, T. Gayden et al. 2017. “Impaired H3k36 Methylation Defines a Subset of Head and Neck Squamous Cell Carcinomas.” Nat Genet 49 (2):180–85. 10.1038/ng.3757.

42. Qu, K., K. Chen, H. Wang, X. Li, and Z. Chen. 2022. “Structure of the Nua4 Acetyltransferase Complex Bound to the Nucleosome.” Nature 610 (7932):569–74. 10.1038/s41586-022-05303-x.

43. Rossetto, D., M. Cramet, A. Y. Wang et al. 2014. “Eaf5/7/3 Form a Functionally Independent Nua4 Submodule Linked to Rna Polymerase Ii-Coupled Nucleosome Recycling.” Embo j 33 (12):1397–415. 10.15252/embj.201386433.

44. Sad, K., D. V. Fawwal, C. Y. Jones et al. 2025. “Histone H3e50k Remodels Chromatin to Confer Oncogenic Activity and Support an Emt Phenotype.” NAR Cancer 7 (1):zcaf002. 10.1093/narcan/zcaf002.

45. Sambrook, J., E. F. Fritsch, and T. Maniatis. 1989. Molecular Cloning: A Laboratory Manual. Second ed. Cold Spring Harbor Laboratory Press.

46. Selleck, William, Israël Fortin, Decha Sermwittayawong, Jacques Côté, and Song Tan. 2005. “The Saccharomyces Cerevisiae Piccolo Nua4 Histone Acetyltransferase Complex Requires the Enhancer of Polycomb a Domain and Chromodomain to Acetylate Nucleosomes.” Molecular and cellular biology 25 (13):5535–42.

47. Sheng, P., Z. Chen, J. Wen, C. Tong, J. Wang, and Z. Du. 2025. “Mg149 Suppresses Anaplastic Thyroid Cancer Progression by Inhibition of Lysine Acetyltransferase Kat5-Mediated C-Myc Acetylation.” Bull Cancer 112 (2):122–34. 10.1016/j.bulcan.2024.11.008.

48. Sikorski, R. S., and P. Hieter. 1989. “A System of Shuttle Vectors and Yeast Host Strains Designed for Efficient Manipulation of DNA in Saccharomyces Cerevisiae.” Genetics 122 (1):19–27. 10.1093/genetics/122.1.19.

49. Spangle, J. M., K. M. Dreijerink, A. C. Groner et al. 2016. “Pi3k/Akt Signaling Regulates H3k4 Methylation in Breast Cancer.” Cell Rep 15 (12):2692–704. 10.1016/j.celrep.2016.05.046.

50. Squatrito, M., C. Gorrini, and B. Amati. 2006. “Tip60 in DNA Damage Response and Growth Control: Many Tricks in One Hat.” Trends Cell Biol 16 (9):433–42. 10.1016/j.tcb.2006.07.007.

51. Steunou, A. L., M. Cramet, D. Rossetto et al. 2016. “Combined Action of Histone Reader Modules Regulates Nua4 Local Acetyltransferase Function but Not Its Recruitment on the Genome.” Mol Cell Biol 36 (22):2768–81. 10.1128/mcb.00112-16.

52. Su, W. P., S. H. Hsu, L. C. Chia et al. 2016. “Combined Interactions of Plant Homeodomain and Chromodomain Regulate Nua4 Activity at DNA Double-Strand Breaks.” Genetics 202 (1):77–92. 10.1534/genetics.115.184432.

53. Szerlong, H., K. Hinata, R. Viswanathan, H. Erdjument-Bromage, P. Tempst, and B. R. Cairns. 2008. “The Hsa Domain Binds Nuclear Actin-Related Proteins to Regulate Chromatin-Remodeling Atpases.” Nat Struct Mol Biol 15 (5):469–76. 10.1038/nsmb.1403.

54. Wagner, E. J., and P. B. Carpenter. 2012. “Understanding the Language of Lys36 Methylation at Histone H3.” Nat Rev Mol Cell Biol 13 (2):115–26. 10.1038/nrm3274.

55. Wang, X., S. Ahmad, Z. Zhang, J. Côté, and G. Cai. 2018. “Architecture of the Saccharomyces Cerevisiae Nua4/Tip60 Complex.” Nat Commun 9 (1):1147. 10.1038/s41467-018-03504-5.

56. Yi, C., M. Ma, L. Ran et al. 2012. “Function and Molecular Mechanism of Acetylation in Autophagy Regulation.” Science 336 (6080):474–7. 10.1126/science.1216990.

57. Zhang, X., D. V. Fawwal, J. M. Spangle, A. H. Corbett, and C. Y. Jones. 2023. “Exploring the Molecular Underpinnings of Cancer-Causing Oncohistone Mutants Using Yeast as a Model.” J Fungi (Basel) 9 (12)10.3390/jof9121187.

58. Zhang, Y., C. M. Shan, J. Wang, K. Bao, L. Tong, and S. Jia. 2017. “Molecular Basis for the Role of Oncogenic Histone Mutations in Modulating H3k36 Methylation.” Sci Rep 7:43906. 10.1038/srep43906.

59. Zhao, J. J., Z. Liu, L. Wang, E. Shin, M. F. Loda, and T. M. Roberts. 2005. “The Oncogenic Properties of Mutant P110alpha and P110beta Phosphatidylinositol 3-Kinases in Human Mammary Epithelial Cells.” Proc Natl Acad Sci U S A 102 (51):18443–8. 10.1073/pnas.0508988102.

60. Zukin, S. A., M. R. Marunde, I. K. Popova, K. M. Soczek, E. Nogales, and A. B. Patel. 2022. “Structure and Flexibility of the Yeast Nua4 Histone Acetyltransferase Complex.” Elife 11 10.7554/eLife.81400.

