## Supplemental Figures for "Overexpression of Eaf1, a subunit of the NuA4 lysine acetyltransferase complex, rescues growth defects in the *Saccharomyces cerevisiae* H3K36M oncohistone model via histone H4 tail acetylation"

### Supplementary Figure Legends

**Supplementary Figure 1. Growth on dropout media confirms plasmid expression.** The dropout plates corresponding to Figure 1C confirm plasmid expression.

**Supplementary Figure 2. Plasmid expression is confirmed by growth on dropout media.** The dropout plates corresponding to Figure 2C confirm plasmid expression.

**Supplementary Figure 3. Representative images of colonies and plasmid-expressing cells.** a) A representative set of plates for H4-4K→A cells in 3B, and b) a representative set of plates for H4-4K→R cells in 3B. A subset of the double mutant colonies in S3B were sequenced and found to have reverted to wildtype at both *HHT2* and *HHF2* genes. So, these colonies do not actually represent viable double mutants.

**Supplementary Figure 4. Growth on dropout media confirms plasmid expression.** The dropout plates corresponding to Figure 4E confirm plasmid expression.

**Supplementary Figure 5. Growth on dropout media confirms plasmid expression.** The dropout plates corresponding to Figure 5 confirm plasmid expression.

### Supplementary Figure 1

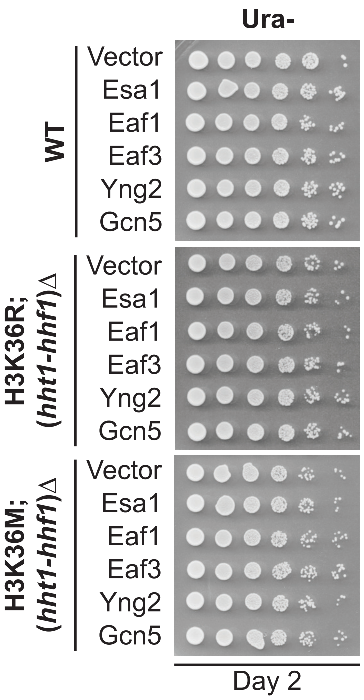

### Supplementary Figure 2

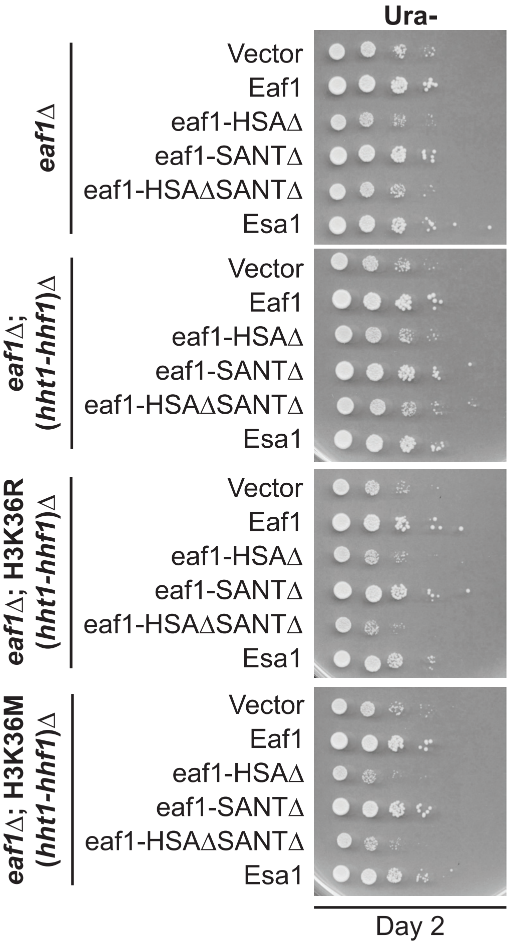

### Supplementary Figure 3

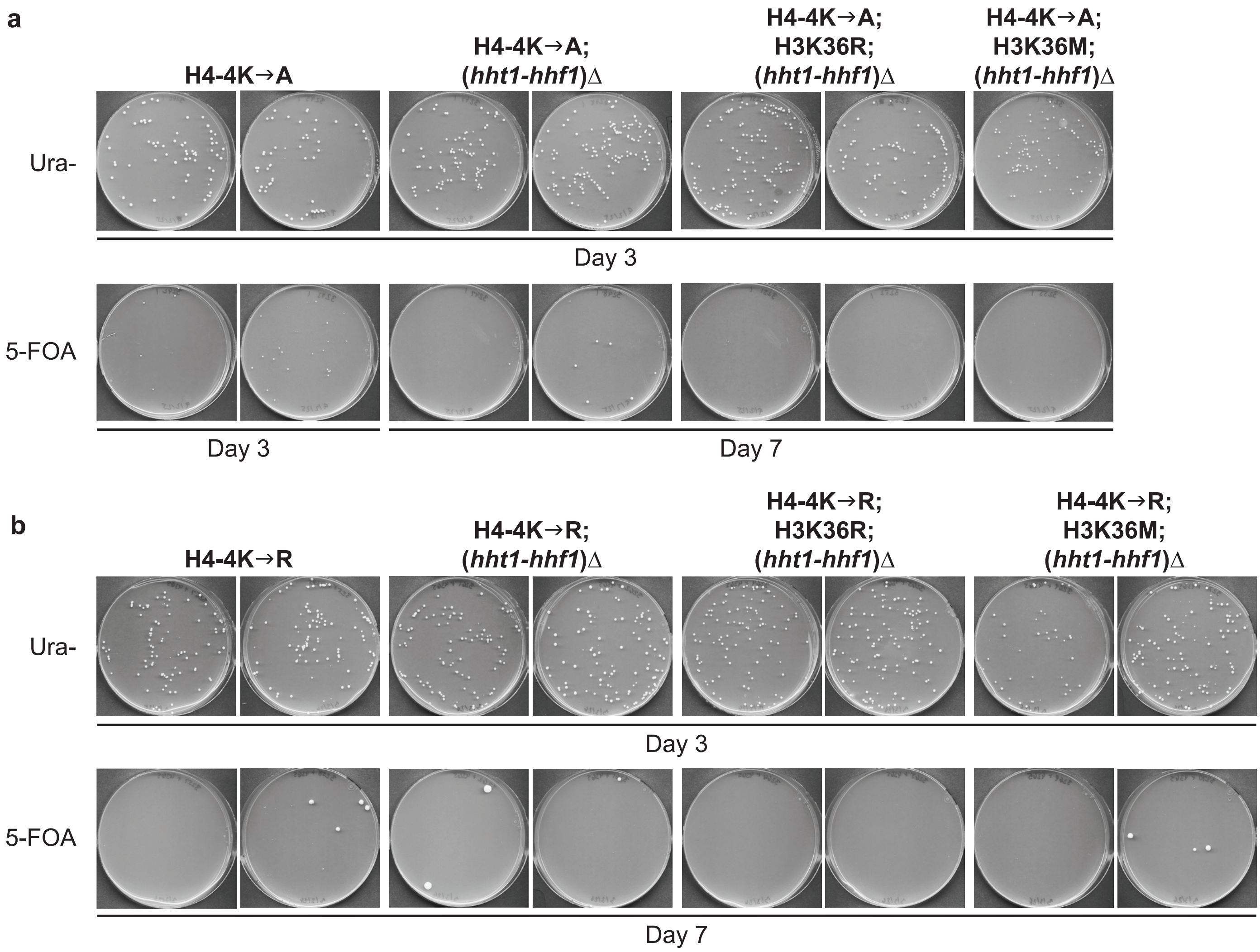

### Supplementary Figure 4

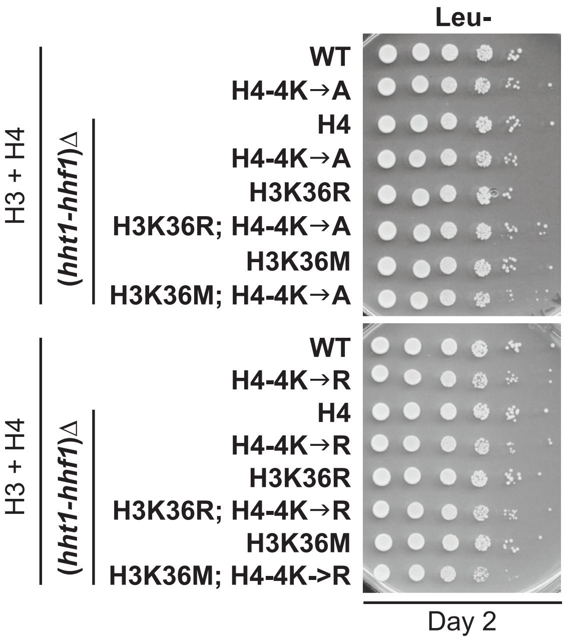

### Supplementary Figure 5

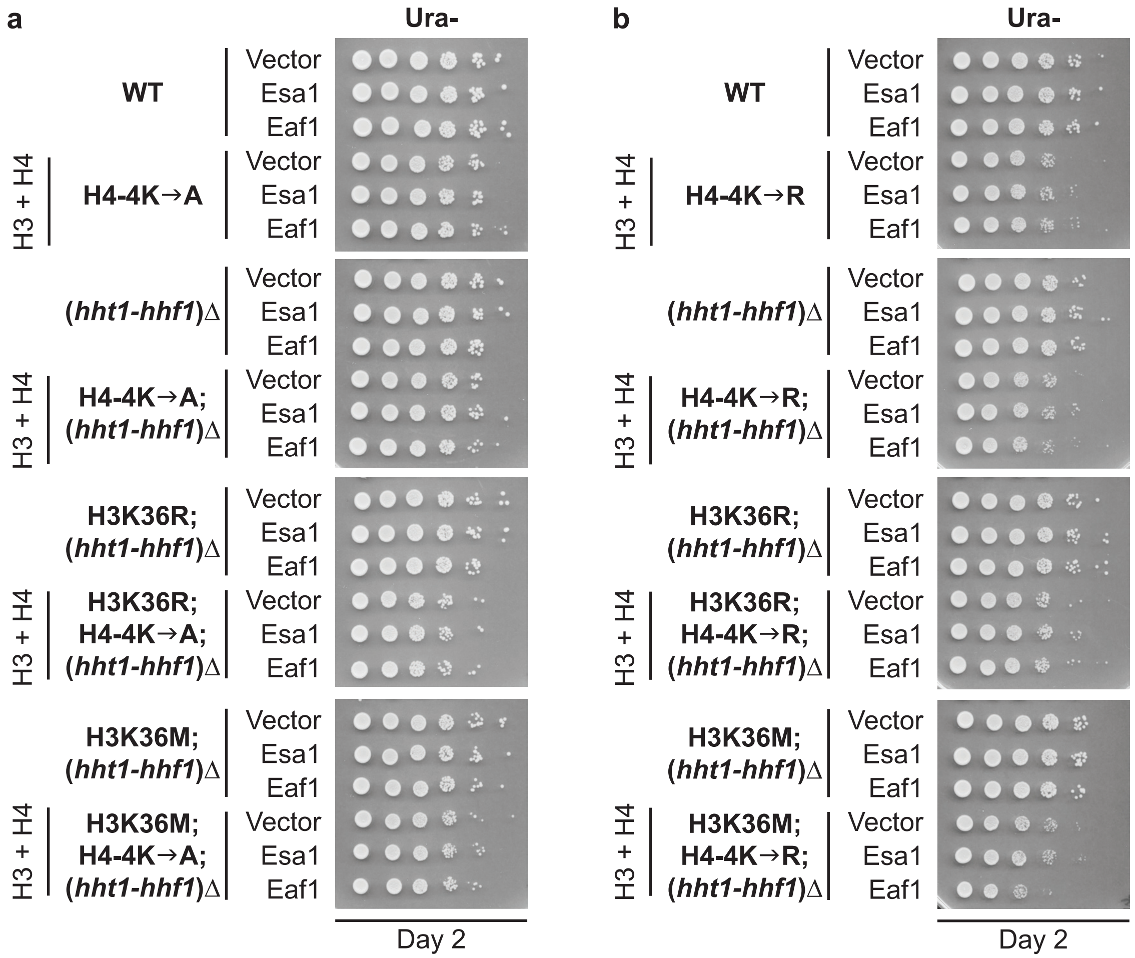

| Strain/Plasmid | Description | Source |
|---|---|---|
| WT (ACY3032) | MATa; *his3*Δ200 *leu2*Δ1 *ura3*-52 *TRP1 lys2*-128delta | This study |
| H3K36R (ACY3088) | MATa; *his3*Δ200 *leu2*Δ1 *ura3*-52 *TRP1 lys2*-128delta *hht2*-K36R (*hht1-hhf1*)Δ::*kanMX* | This study |
| H3K36M (ACY3091) | MATa; *his3*Δ200 *leu2*Δ1 *ura3*-52 *TRP1 lys2*-128delta *hht2*-K36M (*hht1-hhf1*)Δ::*kanMX* | This study |
| *eaf1*Δ (ACY3153) | MATa; *his3*Δ200 *leu2*Δ1 *ura3*-52 *TRP1 lys2*-128delta *eaf1*Δ::*NatMX* | This study |
| *eaf1*Δ; (*hht1-hhf1*)Δ (ACY3157) | MATa; *his3*Δ200 *leu2*Δ1 *ura3*-52 *TRP1 lys2*-128delta (*hht1-hhf1*)Δ::*kanMX eaf1*Δ::*NatMX* | This study |
| *eaf1*Δ; H3K36R; (*hht1-hhf1*)Δ (ACY3159) | MATa; *his3*Δ200 *leu2*Δ1 *ura3*-52 *TRP1 lys2*-128delta *hht2*-K36R (*hht1-hhf1*)Δ::*kanMX eaf1*Δ::*NatMX* | This study |
| *eaf1*Δ; H3K36M; (*hht1-hhf1*)Δ (ACY3165) | MATa; *his3*Δ200 *leu2*Δ1 *ura3*-52 *TRP1 lys2*-128delta *hht2*-K36M (*hht1-hhf1*)Δ::*kanMX eaf1*Δ::*NatMX* | This study |
| H4-4K→A (ACY3244) | MATa; *his3*Δ200 *leu2*Δ1 *ura3*-52 *TRP1 lys2*-128delta *hhf2*-K5A/K8A/K12A/K16A | This study |
| H4-4K→A; (*hht1-hhf1*)Δ (ACY3247) | MATa; *his3*Δ200 *leu2*Δ1 *ura3*-52 *TRP1 lys2*-128delta *hhf2*-K5A/K8A/K12A/K16A (*hht1-hhf1*)Δ + pAC4345 (*HHT2-HHF2*, *LEU2*) | This study |
| H4-4K→A; H3K36R; (*hht1-hhf1*)Δ (ACY3251) | MATa; *his3*Δ200 *leu2*Δ1 *ura3*-52 *TRP1 lys2*-128delta *hhf2*-K5A/K8A/K12A/K16A *hht2*-K36R (*hht1-hhf1*)Δ + pAC4345 (*HHT2-HHF2*, *LEU2*) | This study |
| H4-4K→A; H3K36M; (*hht1-hhf1*)Δ (ACY3255) | MATa; *his3*Δ200 *leu2*Δ1 *ura3*-52 *TRP1 lys2*-128delta *hhf2*-K5A/K8A/K12A/K16A *hht2*-K36M (*hht1-hhf1*)Δ + pAC4345 (*HHT2-HHF2*, *LEU2*) | This study |
| H4-4K→R (ACY3257) | MATa; *his3*Δ200 *leu2*Δ1 *ura3*-52 *TRP1 lys2*-128delta *hhf2*-K5R/K8R/K12R/K16R | This study |
| H4-4K→R; (*hht1-hhf1*)Δ (ACY3261) | MATa; *his3*Δ200 *leu2*Δ1 *ura3*-52 *TRP1 lys2*-128delta *hhf2*-K5R/K8R/K12R/K16R (*hht1-hhf1*)Δ + pAC4345 (*HHT2-HHF2*, *LEU2*) | This study |
| H4-4K→R; H3K36R; (*hht1-hhf1*)Δ (ACY3264) | MATa; *his3*Δ200 *leu2*Δ1 *ura3*-52 *TRP1 lys2*-128delta *hhf2*-K5R/K8R/K12R/K16R *hht2*-K36R (*hht1-hhf1*)Δ + pAC4345 (*HHT2-HHF2*, *LEU2*) | This study |
| H4-4K→R; H3K36M; (*hht1-hhf1*)Δ (ACY3269) | MATa; *his3*Δ200 *leu2*Δ1 *ura3*-52 *TRP1 lys2*-128delta *hhf2*-K5R/K8R/K12R/K16R *hht2*-K36M (*hht1-hhf1*)Δ + pAC4345 (*HHT2-HHF2*, *LEU2*) | This study |
| YEp352 (pAC29) | *URA3*; 2μ; $amp^R$ | Hill et al., 1986 |
| *ESA1* (pAC4190) | *ESA1*; *URA3*; 2μ; $amp^R$ | Lemon et al., 2022 |
| *EAF1* (pAC4400) | *EAF1*; *URA3*; 2μ; $amp^R$ | This study |
| *EAF3* (pAC4401) | *EAF3*; *URA3*; 2μ; $amp^R$ | This study |
| *YNG2* (pAC4402) | *YNG2*; *URA3*; 2μ; $amp^R$ | This study |
| *GCN5* (pAC4293) | *GCN5*; *URA3*; 2μ; $amp^R$ | This study |
| pRS316 (pAC4403) | *URA3*; ARS-CEN; $amp^R$ | Lu et al., 2022 |
| *EAF1* (pAC4404) | *URA3*; ARS-CEN; *EAF1*; $amp^R$ | Lu et al., 2022 |
| *eaf1*-HSAΔ (pAC4405) | *URA3*; ARS-CEN; *eaf1*-HSAΔ (Δ1042–1254); $amp^R$ | Lu et al., 2022 |
| *eaf1*-SANTΔ (pAC4406) | *URA3*; ARS-CEN; *eaf1*-SANTΔ (Δ1936–2121); $amp^R$ | Lu et al., 2022 |
| *eaf1*-HSAΔSANTΔ (pAC4407) | *URA3*; ARS-CEN; *eaf1*-HSAΔSANTΔ (Δ1042–1254, 1936–2121); $amp^R$ | Lu et al., 2022 |
| H3 + H4 (pAC4365) | *HHT2*; *HHF2*; *URA3*; CEN; $amp^R$ | This study |
| H3 + H4 (pAC4345) | *HHT2*; *HHF2*; *LEU2*; CEN; $amp^R$ | This study |
